# Coacervate droplet sequestration of heterogenous nanoplastics with elastin-like polypeptides

**DOI:** 10.64898/2026.03.21.713410

**Authors:** Natalie R. Ling, Aashi Kotecha, Allie C. Obermeyer

**Affiliations:** Department of Chemical Engineering, Columbia University, New York, NY

## Abstract

Nanoplastics generated from plastic waste in our ecosystems are becoming increasingly prevalent as bulk plastics exposed to natural factors like water and sunlight fragment to the nanoscale over time. These incidental nanoplastics span a wide range of physicochemical properties, which makes studying nanoplastic interactions in biological systems difficult. Here, we characterized the behavior of incidental nanoplastics generated through mechanical abrasion within coacervate droplets to probe the surface properties of the nanoplastics. We used elastin-like polypeptides (ELPs) to create hydrophobic or charged coacervate microenvironments. Using optical microscopy and fluorescence quantification, we observed that nanoplastics made from polyethylene terephthalate (nPET), nylon 6 (nPA), and polystyrene (nPS) exhibited distinct partitioning behavior with more favorable interactions with hydrophobic droplets. This indicated that the hydrophobic polymer backbone was the predominate surface feature despite exposed functional groups of the incidental nanoplastics, in contrast to findings with model carboxylated latex nanospheres (nPS-COOH). Furthermore, the selective partitioning of incidental nanoplastics into the hydrophobic droplets was able to capture over 80% of nPET in solution, and after recovery of the protein droplet, was able to cumulatively capture over 75% of the nPET feedstock across multiple cycles. This work explores the nuanced surface characteristics of incidental nanoplastics, expands the application of coacervates as chemical probes, and demonstrates a biopolymer approach for effective nanoplastic removal.

## Introduction

The presence of nanoplastics in the environment is now near ubiquitous^1–9^ and presents a growing threat to both the ecosystem and human health. Formed via the degradation of common commodity plastics, these incidental nanoplastics are a complex mixture of particles that span from millimeters to nanometers.^10^ The nanoscale fragments formed from plastic waste that has leaked into the environment, or incidental nanoplastics, are complex particles with diverse shapes, sizes, and surface chemistries.^11^ Characterizing environmental nanoplastics presents several challenges, as nanoplastics are at low concentrations relative to other materials in the environment and are hard to isolate and identify accurately, limiting characterization primarily to measuring mass concentrations.^12,13^ Due to the broad range of commercial plastics, geographic locations, and routes of degradation, environmental nanoplastics comprise an incredibly diverse range of particles that provide myriad opportunities for interactions with biological systems.^5^ Studying nanoplastic interactions with proteins on a particle-single protein scale reveals adsorbed corona profiles, but is incomplete given current characterization techniques.^14,15^ Therefore, new approaches to probe or control the interactions between environmentally representative nanoplastics and biomolecules are needed.

To address this challenge, we turned to programmable, phase separating biomaterials. Specifically, elastin-like polypeptides (ELPs) are genetically encoded polymers that undergo reversible phase separation, or coacervation, to form protein-rich, liquid-like droplets. Prior work with ELPs has shown that guest residues (Xaa) in the repetitive pentapeptide sequence, Val-Pro-Gly-Xaa-Gly, tune the phase behavior.^16,17^ The guest residue also allows for the chemistry of the coacervates to be precisely controlled at the sequence level,^18^ making them ideal to probe the selectivity of particle recognition. ELPs can form simple coacervates dominated by hydrophobic interactions with a canonical Val in the guest residue position.^19^ Or, when functionalized with charged residues, such as Lys and Glu, ELPs can form complex coacervates driven by electrostatic interactions.^20^ These two systems act as complementary chemical probes; while bulk plastics are hydrophobic, nanoplastics have also been shown to interact with polyelectrolyte complexes.^21^ This allows us to systematically investigate the collective surface properties of nanoplastics via optical microscopy by observing nanoplastic partitioning behavior into distinct protein-rich environments. We hypothesized that this approach would not only reveal the key physicochemical drivers of nanoplastic-protein interactions but also uncover a potential strategy for a broad approach to sequestration.

Here, we created incidental nanoplastics of common plastics and investigated the partitioning of these nanoplastics in two complementary protein coacervates, exploring a hydrophobic and a charged microenvironment. We showed that nanoplastics formed from stirring bulk plastics in water at ambient conditions were diverse in size and shape. Critically, the incidental nanoplastics also had varying surface characteristics dependent on the polymer backbone. We evaluated how nanoplastics from three common plastics, polyethylene terephthalate (PET), polycaprolactam (nylon 6, PA), and polystyrene (PS), interacted with polypeptides *in vitro* by observing their partitioning behavior within coacervates and probed the features driving nanoplastic interactions with these polypeptides. We observed highly favorable interactions with the hydrophobic, simple coacervates regardless of plastic type, while interactions with charged, complex coacervates had a stronger dependence on the nanoplastic composition. We further showed that the behavior of the mechanically generated nanoplastics was distinct from a commonly used model latex nanoplastic.^22–25^ Leveraging this fundamental insight, we successfully applied the hydrophobic ELP coacervate as a recyclable biosorbent for the efficient removal of nanoplastics from water. This work establishes polypeptide coacervates as a powerful platform for both interrogating the interfacial properties of nanoplastic pollutants and developing new, biologically inspired remediation technologies.

## Results and Discussion

### Mechanically generated nanoplastics from common plastics are diverse in morphology

Several model nanoparticles have been developed to probe the impact of nanoplastics on living systems.^26^ Commonly studied primary, or intentionally created, nanoplastics include commercially available nanoparticles^22– 24,27,25^ or nanoplastics formed from dropwise reprecipitation of dissolved commodity plastics at the lab scale.^28,29^ Model secondary, or fragmentation driven, nanoplastics are typically generated by cryomilling or sequentially ball milling pristine pellets or consumer plastic products.^30,15,31^ These models have provided a basic foundation for exploring the effect of nanoplastics on biological systems, but are unable to capture the nuanced characteristics of the nanoplastics in our ecosystems.^32^ At the nanoscale, the surface area to volume is higher, which exaggerates the role of surface characteristics in nanoplastic behavior compared to plastics at the bulk scale. Environmental nanoplastics are secondary nanoplastics that form from bulk plastics exposed to water, sunlight, and living organisms in natural ecosystems. This leads to mechanical, chemical, and biological fragmentation that forms micro and nanoplastics with diverse morphologies and surface chemistries.^5,11^

To better mimic environmental nanoplastics, we devised a method to generate, isolate, and fluorescently stain nanoplastics that reflect the natural mechanical abrasion process, as mechanical degradation is applicable to all types of plastics. We selected PET, PA, and PS and used pristine 3 mm plastic granules to make nanoplastics through the collision of bulk plastics in an aqueous environment. We stirred the plastic granules in ultrapure water (UPW) at ambient conditions for 1 month and through a series of centrifugation steps, isolated a solution of nanoplastics (SI Fig. 1). To visualize and quantify the nanoplastics, we adapted a method to label the nanoplastics with Nile red^33^ without observably altering the nanoplastics. Both unaltered and labeled nanoplastics were characterized by scanning electron microscopy (SEM) for size and morphology, dynamic light scattering (DLS) for size and zeta potential, and nanoparticle tracking analysis (NTA) for size and concentration. No discernable physical differences were observed between the non-fluorescent and fluorescent nanoplastics (SI Fig. 3–5). We also characterized commercially available red fluorescent 0.1 μm carboxylate-modified latex primary nanoplastics (nPS-COOH) to compare with the ambiently generated Nile red labeled PET nanoplastics (nPET), nylon nanoplastics (nPA), and polystyrene nanoplastics (nPS).

The stresses induced by stirring bulk plastics in water produced discernable, nonuniform nanoplastics in the 10-300 nm size range for all the plastics tested. The size distribution of nPET and nPA by DLS and NTA showed particles with monomodal intensity and number size distributions of 100–300 nm (Fig. 1e,f). SEM micrographs confirmed the given size range but showed significant heterogeneity in shape and morphology (Fig. 1a,b). While a majority of the nPET were rounded but non-spherical (Fig. 1a), stick-like shapes and other small rough clusters of nPET were also observed via SEM (SI Fig. 5a). The morphology of nPA varied more significantly, with some nPA fragments forming web-like and sharp structures (SI Fig. 5b). In contrast, the size distribution for nPS was bimodal with nanoplastics both below 10 nm and above 200 nm in size (Fig. 1g), which was also seen in SEM images (Fig. 1c). We added a nonionic surfactant for nPS size distribution measurements via DLS to stabilize the extremely hydrophobic nPS in solution. Lastly, the commercial nPS-COOH were confirmed to be 100 nm in size via DLS and NTA with monomodal distributions (Fig. 1h). The nPS-COOH were also near-perfect spheres (Fig. 1d), highlighting the physical discrepancies between commercially available nanoplastics and the nanoplastics generated through an environmentally representative process. In addition to size distributions, NTA also provided concentration approximations, with the stock nanoplastic samples after Nile red staining ranging from 10^7^–10^9^ particles/mL (approximately 30–3,000 ng/mL, SI Table 1), which was sufficiently high in concentration to mimic environmental concentrations of 1.5–32 ng nanoplastics/mL.^7^

**Figure 1.**
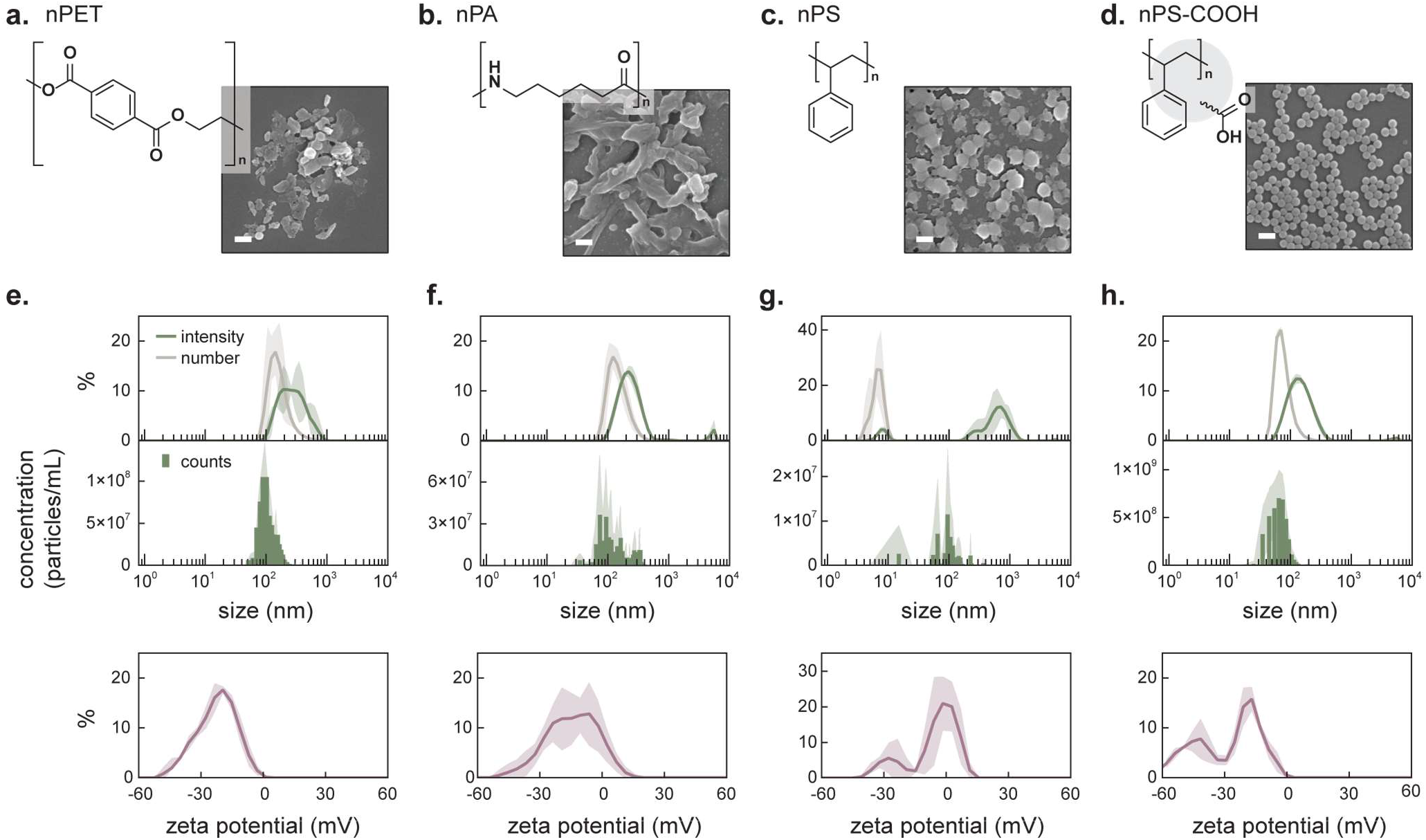
Characterization of environmentally representative nanoplastics. SEM images of nanoplastics generated through stirring of **a**. PET, **b**. nylon 6 (PA), and **c**. PS in water at ambient conditions for 1 month, and **d**. 0.1 μm carboxylate-modified latex microspheres (nPS-COOH). Micrographs are representative of the majority of nanoplastics seen across several batches of nanoplastic production. Scale bars, 200 nm. Particle size distributions obtained by DLS (top) and NTA (middle), and zeta potential (bottom) distributions obtained by DLS for **e**. PET nanoplastics (nPET), **f**. nylon 6 nanoplastics (nPA), **g**. polystyrene nanoplastics (nPS), and **h**. carboxylated latex nanoparticles (nPS-COOH). For size and zeta potential DLS data, the plots show the mean value and standard deviation (shaded) from three measurements of the same nanoplastic stock solution. For NTA data, the histograms represent the mean and standard deviation (shaded) of three technical replicates of the same nanoplastic stock solution.

While size and shape analysis indicated minor differences between plastic types and distinct differences compared to primary nanoplastics, zeta potential measurements highlighted additional discrepancies between the surface properties of the mechanically degraded nanoplastic samples. For nPET and nPA, an overall negative zeta potential was observed (Fig. 1e,f) for the stock solutions in water, and these samples were colloidally stable, after resuspension by water bath sonication, for several days in water and at least one day in tris buffer with varying salt concentrations (SI Fig. 6). While primary nylon nanoplastics reported positive zeta potentials,^29^ likely from the amide group exposed after reprecipitation of complete polymer chains, we hypothesize that the observed negative charge in these mechanically generated nanoplastics was due to the fragmentation of nPA in an aqueous solution. Hydrolysis of the amide group in the polymer backbone would lead to both an exposed amine and carboxylic acid group,^34^ resulting in a negative zeta potential at pH 7. For nPS, bimodal peaks were also observed in the zeta potential with the primary population of nPS showing a neutral zeta potential (Fig. 1g). This deviated from the two dominant, highly negative zeta potentials of nPS-COOH (Fig 1h), which can be attributed to the hydrophilic, anionic polymer coating of the commercial nanospheres. Given the different surface properties of these nanoplastics indicated by the zeta potential, we hypothesized that they may have differential interactions with the two coacervate microenvironments. Zeta potential analyses are fundamentally limited by the measurement methods,^35^ so with an approximate understanding of the size and surface properties of the mechanically generated nanoplastics, we next created distinct protein rich environments using phase-separated polypeptides to probe nanoplastic partitioning and predominant surface properties on a holistic scale.

### Simple and complex ELP coacervates form low volume dense phases modulated by salt concentration

Given the distinct microenvironments possible for phase-separated droplets, we evaluated how nanoplastics interacted with protein coacervates as biologically relevant scenarios to probe nanoplastic properties. We hypothesized that the coacervate composition would affect how a given nanoplastic interacts with the droplet, as previous research has shown that client-scaffold interaction strength determined if and where a large particle partitioned in a droplet.^36^ We used elastin-like polypeptides (ELPs), a synthetic polypeptide with the repeat sequence [VPGXG]_n_ based on tropoelastin, to create protein droplets. The phase behavior of ELPs is well-characterized and modifiable via repeat length (n) and identity of the guest residue (X).^16,37^ To create droplets with a hydrophobic microenvironment, we chose ELP-V150, or [VPG<u>**V**</u>G]_150_, as it can form simple coacervates with nonspecific hydrophobic interactions driven by the exclusion of water and salt (Fig. 2a).^38^ ELP-V150 has a lower critical transition temperature (LCST) that can be decreased with increasing salt concentration or increasing chain length.^17,19^ To create an alternate, more hydrophilic microenvironment, we used modified ELPs with lysine (ELP-K30, [VPG<u>**K**</u>G]_30_) and aspartic/glutamic acid (ELP-DE30, or [VPG<u>**D**</u>GVPG<u>**E**</u>G]_15_) guest residues (SI Table 2, SI Fig. 7) to introduce positive and negative charges throughout the polypeptide to facilitate electrostatically driven complex coacervation (Fig. 2d).^20^ We mapped the phase behavior of ELP-V150 simple coacervates and ELP-K30, ELP-DE30 complex coacervates as a function of salt (Fig. 2b,e) up to 1 M NaCl. We coupled salt dependence with temperature to probe the LCST properties of ELP simple coacervates, and the positive charge fraction (*f*+), calculated as [ELP-K30]/[total ELP], to evaluate the tolerance for charge imbalance of ELP complex coacervates. The broad range of conditions that promote phase separation give the coacervate system flexibility to accommodate a wide range of solution and nanoplastic properties.

**Figure 2.**
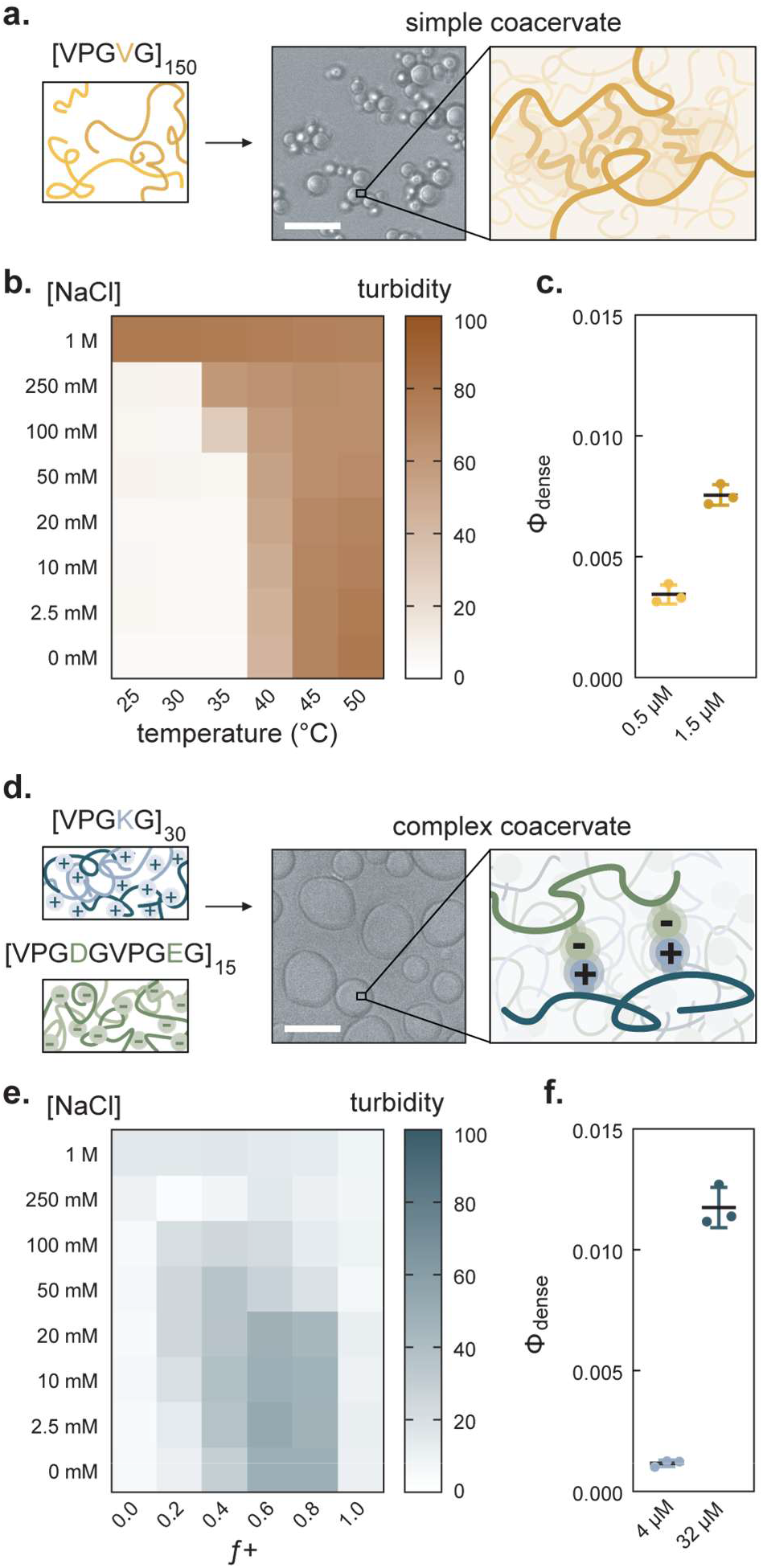
Phase behavior of ELP simple and complex coacervates. Schematic and representative brightfield microscopy image of coacervate formation for **a.** ELP-V150 at high salt conditions driven by hydrophobic interactions and **d**. ELP-K30 with ELP-DE30 at low salt conditions driven by electrostatic interactions between lysine and aspartic/glutamic acid side chains. Scale bars, 10 μm. Phase diagram for **b**. simple coacervates of 1.5 μM ELP-V150 varying temperature and salt concentration and **e**. complex coacervates of 32 μM total of ELP-K30, ELP-DE30 varying the positive charge fraction and salt concentration. Measured volume fraction of c. the simple coacervate dense phase at 0.5 μM (0.03 mg/mL) and 1.5 μM (0.09 mg/mL) ELP-V150 and **f**. the complex coacervate dense phase at total ELP concentration of 4 μM (0.06 mg/mL) and 32 μM (0.50 mg/mL) ELP-K30, ELP-DE30. Turbidity heat map values represent the mean of n=3 technical triplicates. Volume fraction data points represent individual replicates (n=3), black bar represents the mean, and error bars represent the standard deviation.

With a relevant phase space mapped, we selected the optimal solution conditions for droplet formation to evaluate nanoparticle partitioning. We observed that simple coacervation occurs with ELP-V150 at room temperature with elevated salt concentrations of 1 M NaCl, and droplet formation occurred above 40 °C regardless of salt concentration (Fig. 2b). Furthermore, by using temperature to control phase separation, the system demonstrated reversible droplet formation for at least five cycles at 20 mM NaCl while maintaining a transition temperature of 35.1 ± 0.2 °C (SI Fig. 8). For simplicity, all subsequent experiments studied simple coacervates at room temperature with 1 M NaCl. For the complex coacervates, we observed a minor charge imbalance for optimal complex coacervate formation, possibly due to residual impurities following purification and the lower precision of protein concentration determination for atypical or low sequence complexity polypeptides.^39^ The most turbid samples were seen at *f*+ 0.6 and salt concentrations below 20 mM NaCl (Fig. 2e). These micron-scale, protein-rich droplets formed via phase separation gravitationally settle over a few hours, as brightfield microscopy showed the simple coacervate formed spherical droplets around 2–5 μm in diameter and the complex coacervates wetted the surface and formed wider puddles spanning 4–12 μm. Over time, the droplets in solution accumulated at the bottom of the vessel, creating a distinct dilute, protein-poor phase and the accompanying small, protein-rich droplets.

The coacervate droplets make up a minimal fraction of the solution volume and can be collected by centrifugation to create a macroscale droplet at low input protein concentrations. Imaging of the centrifuged droplet allowed estimation of the dense phase volume fraction (ϕ_dense_) (SI Fig. 9). Simple coacervates prepared at 1.5 μM total ELP-V150 (0.09 mg/mL), occupied less than 0.8% of the total solution volume (Fig. 2c) and the complex coacervates, at 32 μM total ELP-K30, ELP-DE30 (0.50 mg/mL) occupied less than 1.4% of the total solution volume. With consideration to limit the protein material required to capture nanoplastics, we aimed to study the nanoplastic-coacervate system at a protein concentration close to the saturation concentration. While droplets did form at concentrations as low as 0.1 μM ELP-V150 and 1 μM ELP-K30, ELP-DE30 (SI Fig. 11), we chose 0.5 μM total ELP-V150 (0.03 mg/mL) and 4 μM total ELP-K30, ELP-DE30 (0.06 mg/mL, *f*+ 0.6) at 1 M NaCl and 20 mM NaCl, respectively (Fig. 2a,d), as the optimal concentration for all following experiments. This balanced the minimal use of protein with consistent turbidity upon mixing and abundant, settled droplets on the time scale of 30 min to 2 h (SI Fig. 11,12). At these reduced concentrations, the dense phase volume fraction of the droplets decreased to less than 0.4%. We note that in addition to different microenvironments, properties like droplet water content and density also differ for the two types of coacervates, ultimately impacting the experimental volume fractions. To understand nanoplastic-coacervate interactions, we focused on quantifying the selective partitioning of nanoplastics into the low volume dense phase. Using these measured volume fractions, the nanoplastic-coacervate systems could result in approximately 200-to 500-fold concentration of the nanoplastics within the droplets.

### Nanoplastics of select commodity polymers show differential partitioning in coacervates

After characterizing the nanoplastics and coacervates independently, we next sought to evaluate how the nanoplastics interacted with the two droplet systems. To observe the partitioning behavior of nPET, nPA, and the carboxy-nPS within droplets, we started with lower concentrations of nanoplastics at 100 ng/mL, nearing the magnitude of reported environmental samples while maintaining enough particles per frame of view.^7^ We also then looked at nanoplastic partitioning up to 2.5 μg/mL to more readily visualize and quantify partitioning behavior within droplets. The concentration of nPS was instead evaluated at lower concentrations, ranging from 4 ng/mL to 30 ng/mL, since the low density and hydrophobicity of nPS limited the maximum concentration of the prepared solution. The nPS concentration gradient was chosen to cover one order of magnitude visible via fluorescence microscopy. By observing the formation of ELP coacervates in various nanoplastic containing solutions, we gained insight into the surface interactions of nanoplastics that primarily determine nanoplastic partitioning behavior. We have also shown that in addition to partitioning biomacromolecules, coacervates also readily partition a range of nanoplastics with compatible surface chemistries, similar to what has been observed for uniform nanoparticles used for microrheology.^36^

Though PET and nylon have fundamentally different polymer backbones, with PET containing ester functional groups and aromatic features while nylon has amide groups and aliphatic functionality, nPET and nPA exhibited similar partitioning behavior with the ELP coacervates. Both appeared to partition within the simple coacervates but localized at the interface of the complex coacervates (Fig. 3b,c). This suggested that though both nPET and nPA may have exposed charged functional groups, the main contributing interactions on the nanoplastic surface were from the hydrophobic polymer backbone. These dominating hydrophobic interactions led to nanoplastic inclusion within the simple, hydrophobic droplets, while the milder affinity between the charged functional groups and the electrostatic-driven droplets led to the localization of the nanoplastics along the interface. While the complex coacervates provided an overall net neutral environment, the droplet itself has fleeting, dynamic electrostatic interactions and a higher charge density that may not be as favorable for the low charge density nanoplastic particles.^40^

**Figure 3.**
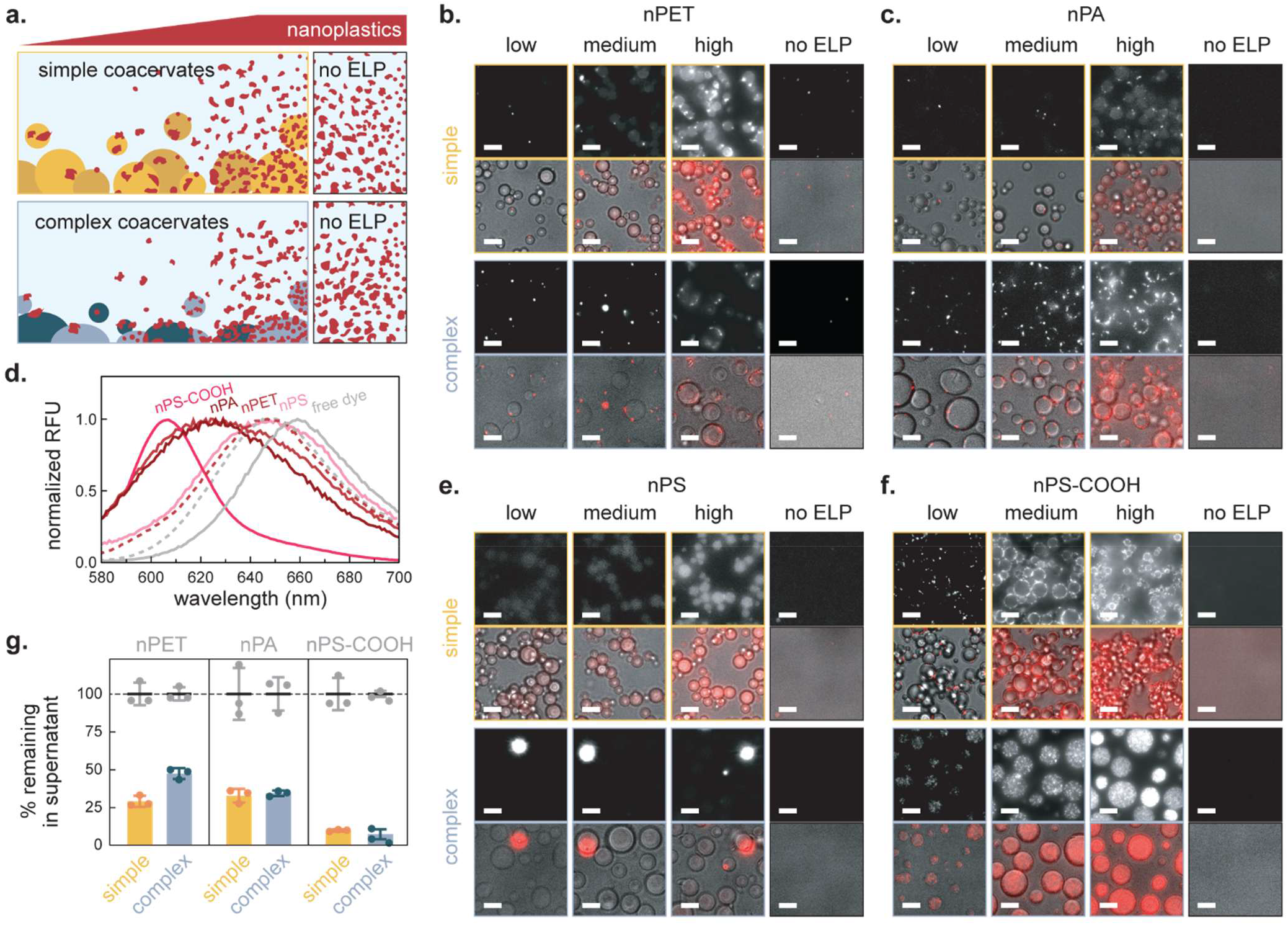
Partitioning behavior of common nanoplastics in ELP coacervates. **a.** Schematic illustrating the experimental conditions for observing nanoplastic interactions with both simple and complex coacervates with increasing nanoplastics; concentrations span environmentally relevant to highly concentrated. For **b.** nPET, **c.** nPA, and **f.** nPS-COOH, the nanoplastic concentrations are 100 ng/mL, 500 ng/mL, and 2.5 pg/mL, with the no ELP control at 2.5 pg/mL. For **e.** nPS, the concentrations are 3.8 ng/mL, 7.5 ng/mL, and 30 ng/mL, with the no ELP control at 30 ng/mL. All nanoplastic concentrations are approximations determined using the average particles/mL count and mean size from NTA with the simplifying assumption that all nanoplastics are spheres and have the same density as the bulk plastic. Simple coacervates (top) are formed via addition of 0.5 μM ELP-V150 into a 1 M NaCI, 10 mM tris, pH 8.0 solution of the respective nanoplastics, and complex coacervates (bottom) are formed via addition of 2.4 μM ELP-K30 and 1.6 μM ELP-DE30 into a 20 mM NaCI, 10 mM tris, pH 8.0 solution of the respective nanoplastics. Widefield mCherry fluorescence (λ_ex_ = 555 nm) and DIC microscopy were used to image nanoplastic partitioning after 4 h. Scale bars, 5 μm. **d.** Normalized fluorescence emission spectra for free Nile Red dye and fluorescently-labeled nanoplastics in water (solid lines) and in 50% ethanol (dashed line) (λ_ex_ = 560 nm for nPET, nPA, and free dye; 540 nm for nPS; 580 nm for nPS-COOH). **g.** Fluorescence quantification of free nanoplastics suspended in solution 1 h after simple and complex coacervate formation, or the amount not partitioned into droplets. Droplet settling was accelerated with centrifugation at 1,000*xg* for 10 min, and nanoplastic concentrations were normalized to the no ELP control conditions as some was lost due to surface adsorption (initial input concentrations of 100 ng/mL nPET, 1 pg/mL nPA, 1 pg/mL nPS-COOH). nPS was not quantified due to insufficient fluorescence intensity. Data are presented as the technical replicates (n=3) for each plastic and coacervate set, bars and black lines are the mean, and error bars indicate the standard deviation.

When comparing the two types of polystyrene nanoplastics, we observed near opposite behavior between the incidental nPS and the commercially manufactured nPS-COOH. The nPS were included within the simple coacervates in a similar way to the nPET and nPA but aggregated in the presence of the complex coacervates (Fig. 3e). The carboxy-nPS, however, localized at the interface of the simple coacervates, while fully internalizing into the complex coacervates (Fig. 3f). Furthermore, the carboxy-nPS, often used for microrheology,^41^ within the hydrophilic droplets exhibited Brownian motion and maintained mobility within the droplets during the 4 h observation period. The behavior of the nPS and carboxy-nPS highlighted the clear connection between the charged surface of nanoplastics and how they interacted with droplet environments relative to the overall hydrophobicity. Even when comparing the highly charged carboxy-nPS to the incidental nPET and nPA, all of which share a negative zeta potential distribution peaking around -20 mV, the behavior of the carboxy-nPS in these biological systems is drastically different.

Next, we reasoned that if the nanoplastics favorably interacted with the droplets, then the droplet could serve as an encapsulation method for concentrating and extracting nanoplastics. After forming droplets within nanoplastic solutions and collecting a bulk droplet, we measured both the concentration of fluorescent nanoplastics remaining suspended in solution and the amount of nanoplastics sequestered by the droplets. We found that given the solvatochromaticity of Nile red, the three mechanically generated nanoplastics had distinct fluorescence spectra (Fig. 3d). We were able to enhance the fluorescence signal of nPET by adding an equal volume of ethanol immediately prior to measurement, which allowed us to reliably quantify nPET concentrations down to several nanograms per milliliter (SI Fig. 13). There was insufficient signal from nPS at the maximum possible concentration (100 ng/mL), so unfortunately, quantification of nPS partitioning was not possible. Across all plastics, we observed that the droplets facilitated the capture of a majority of the nanoplastics after 1 h (Fig. 3g), as measured by the concentration of nanoplastics remaining suspended in solution. We also redissolved the droplet to quantify the captured nanoplastics. Using the error percent of the mass balance, we observed that there is variable loss of nPA and nPS-COOH within the experimental setup (SI Fig. 15), likely due to surface adsorption to the well plates and higher nPA and nPS-COOH concentrations leading to an incomplete resuspension of the pelleted nanoplastics.

After coalescing the droplets into a bulk macrophase droplet to create distinct dense and dilute phases, we found that the overall amount of captured nanoplastics between the two droplet types was similar, implying that the location of the nanoplastics in the droplet as observed by microscopy was irrelevant for the removal efficiency. Furthermore, while ∼50–70% of the nPET and nPA were removed, almost 90% of the nPS-COOH nanoplastics were captured by both droplets. This highlighted the distinction between more representative nanoplastics and model plastic nanoparticles, as the surface characteristics dominate bulk material identity for determining how nanoplastics interact with surrounding biological environments or potential sorbent materials.

### Partitioning of nPET within droplets is thermodynamically favorable

With a base understanding of how nanoplastics interact within simple and complex coacervates, we then explored the driving force for nanoplastic incorporation into these droplets (Fig. 4a). All prior experiments mimicked a realistic mixing order, as soluble polypeptides were added into dilute nanoplastic solutions at the appropriate conditions for phase separation. Given the prevalence of PET in environmental waste^42,43^ and the low limit of nPET detection in our assays, we used nPET as the example nanoplastic in subsequent studies. We observed that as droplets settled with gravity, they brought the partitioned nanoplastics with them for both simple and complex coacervates (Fig. 4b,c). With the rapid formation of droplets, nanoplastics could be trapped in a metastable state within the protein-dense droplet and the apparent nanoplastic capture over the observable period could be an artifact of diffusion limitations. However, if the partitioning of the nanoplastics in the droplets was thermodynamically favorable, then we would expect the nanoplastics to remain encapsulated within the droplets. While the mechanism driving nanoplastic partitioning is not critical for applying coacervates as a concentration method, it provides a better understanding of coacervate dynamic properties.

**Figure 4.**
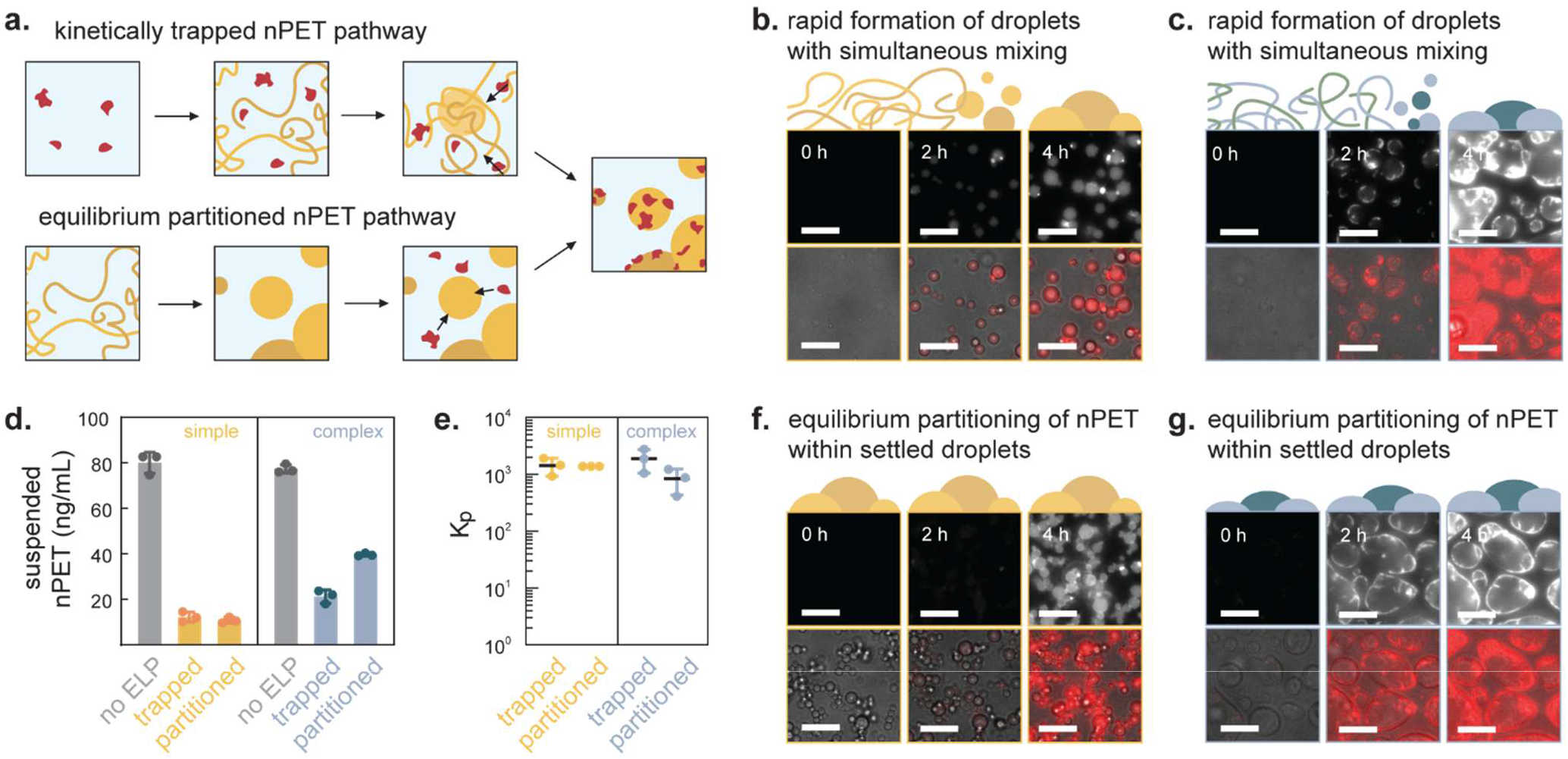
Thermodynamic control of nPET partitioning in simple and complex coacervates. **a**. Schematic of experimental mixing order to determine the driving force for nanoplastic partitioning within droplets. The top pathway illustrates simultaneous mixing of the nPET and polypeptide so nanoplastics are present during droplet formation and coalsecence for **b.** simple coacervates and **c.** complex coacervates. The bottom pathway uses **f.** simple or **g.** complex coacervate droplets that have settled that are then exposed to nanoplastics. Scale bars, 10 μm. Fluorescence quantification of **d.** nPET suspended in bulk solution 2 h after nanoplastic addition and the **e.** estimated partition coefficients of nPET within the simple and complex coacervates for both mixing orders. Bars and black lines represent the mean. Error bars represent the standard deviation with n=3 technical replicates.

By rearranging the mixing order of the nanoplastics and proteins, we confirmed that the localization of nPET within or around the droplet is due to the nPET affinity for the simple and complex coacervate environment. To confirm nPET encapsulation within a droplet as the equilibrium state, we first let simple and complex coacervates form and settle for 2 h in the absence of nanoplastics. We then gently added in a solution of nPET and observed via microscopy that they eventually partitioned within or at the interface of the preformed droplets over the course of 4 h (Fig. 4f,g). The observed nPET partitioning within existing droplets suggested that these entangled ELP scaffolds are highly dynamic^44^ and can internally rearrange to accommodate the client particles, provided the affinity is sufficiently strong.^36^

For the simple coacervates, more than 80% of the nanoplastics were collected by the droplets with no difference between the simultaneous mixing of all components and the addition of nPET to preexisting droplets (Fig. 4d). For the complex coacervates, we observed that slightly fewer were removed when the nPET had to diffuse and partition within the settled droplets as opposed to starting with a homogenous nPET-protein mixture. We attributed this to a slower rate of nPET partitioning reflecting the weaker interaction strength between the nPET and complex coacervates. Partitioning was quantified 2 h after the addition of nPET to minimize plastic aggregation and adsorption (SI Fig. 18,19) while maximizing time for equilibration with the droplets. Based on the previously measured volume fraction of the simple (ϕ_dense_ ≈ 0.004) and complex (ϕ_dense_ ≈ 0.001) coacervates, we approximated a nPET partition coefficient on the order of 10^3^ for both types of coacervates.

### Droplets can be recovered and reused to remove nanoplastics over multiple cycles

Lastly, to demonstrate that biopolymer coacervates can enrich nanoplastics as a potential removal method, we explored the use and reuse of the simple coacervate to sequester and accumulate nanoplastics. Since coacervate formation is reversible and the polypeptide in its two-phase state is easily recoverable, we expected that upon adding a given amount of ELP to a nanoplastic solution, droplets would form and concentrate nanoplastics within a small dense phase, and subsequently, the dilute, largely nanoplastic-free phase could be extracted. Then, depending on the system, the polypeptides in the enriched dense phase could be resolubilized either in a cold solution or a high salt solution and reused for another round of nanoplastic capture (Fig. 5a). We used simple coacervates to demonstrate this concept, as the polypeptide can be purified with high yields, the hydrophobicity of the polypeptide results in a more universal capture of common nanoplastics, and the LCST behavior allows for flexible manipulation of droplet formation as a function of salinity or temperature.

**Figure 5.**
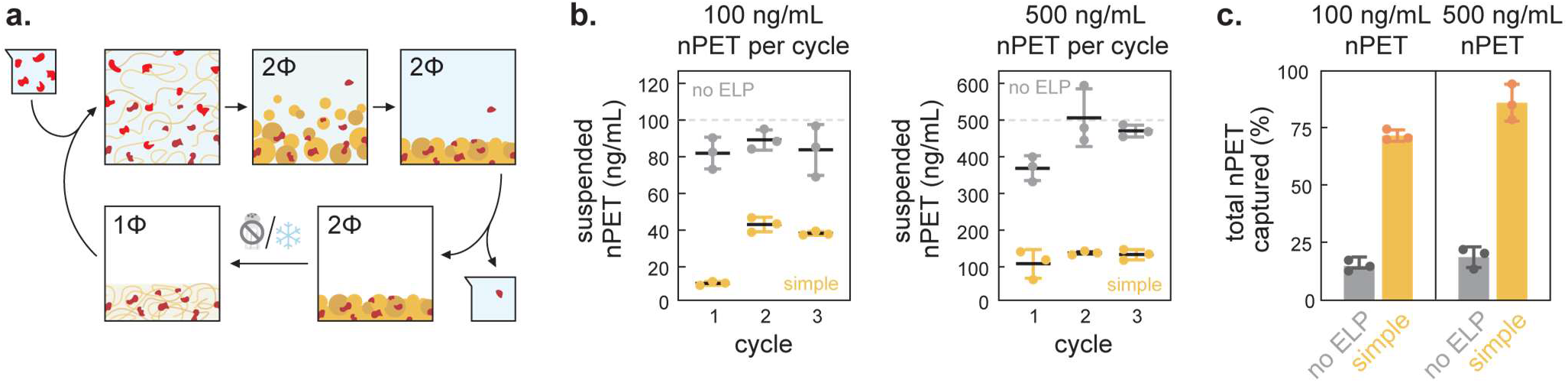
Resuability of simple coacervates for nanoplastic removal. **a**. Schematic depicting the batch processing of dilute nanoplastic feeds to concentrate nPET within droplets over several cycles, **b.** Using an initial input of 0.5 μM ELP-V150 in 1 M NaCI at room temperature, the droplets were isolated, resuspended, and repartitioned through three cycles with fresh 100 ng/mL nPET and 500 ng/mL nPET feed solutions, **c.** The total removal efficiency of nanoplastics after three cycles of droplet reuse from measuring the nanoplastics accumulated in the dense phase. Lines and error bars represent the mean and standard deviation with n=3 technical replicates shown as individual data points.

Using one initial input of 0.5 μM ELP-V150 over three cycles at room temperature with 1 M NaCl, we removed a majority of the nanoplastics each cycle for input particle feedstocks of 100 ng/mL or 500 ng/mL nPET (Fig. 5b). The per cycle efficiency dropped after the first cycle, likely due to loss of some ELP-V150 with the dilute phase removal. We also observed that at the higher nPET concentration of 500 ng/mL, there was a higher removal efficiency cycle over cycle. We attributed this to denser droplets from greater nanoplastic loading at the elevated nPET stock concentration, which led to improved settling and isolation of the dense phase. Totaling across three cycles of droplet reuse, we were able to capture over 70% of nPET at more environmentally relevant concentrations of 100 ng/mL and almost 90% of the nPET at elevated nanoplastic concentrations of 500 ng/mL (Fig. 5c). In this experiment, we used high salt conditions at room temperature as an example case for nanoplastic removal. However, since the onset of phase separation for these simple coacervates can be manipulated in tandem with salt concentration and temperature, freshwater feeds could be heated to above 40 °C, while saltwater feeds could be processed at room temperature. Temperature then becomes a complementary tool to accommodate the varying salinity across water sources for droplet formation and nanoplastic partitioning.

## Conclusion

In this work, we characterized morphologically heterogenous incidental PET, PA, and PS nanoplastics. Though the mechanically generated nanoplastics are made in highly controlled settings using pristine plastics, the protocol for creating and isolating the nanoplastics can be easily applied to most bulk plastics to improve nanoplastic models. To better understand the physicochemical diversity of incidental nanoplastics, we observed how the nPET, nPA, and nPS interacted with a hydrophobic and hydrophilic microenvironment formed through simple and complex coacervation of modified ELPs. Overall, we saw that the hydrophobic, simple coacervate showed more favorable and universal partitioning of the mechanically generated nanoplastics. This indicated that even with exposed polymer functional groups, the hydrophobic nature of nanoplastics was the predominant feature. We also compared the behavior of the mechanically generated nanoplastics with a model nanoparticle, nPS-COOH, and found distinct differences in behavior between the surface modified latex spheres and the incidental nanoplastics. Here, we show that using more representative nanoplastics is critical for understanding potential downstream biological effects. At the nanoscale, as this work has shown, nanoplastic properties beyond the polymer composition can impact how they interact with biological systems.

Given that nanoplastics selectively interact with protein coacervate droplets, we showed that coacervation can be utilized to concentrate and remove nanoplastics with polypeptides. The droplets could readily be recovered for repeated nanoplastic capture. More fundamentally, our results reinforce that protein coacervates are highly dynamic and can rearrange and accommodate select 10-300 nm nanoplastic clients. While only two distinct coacervate microenvironments, one hydrophobic and one charged, were explored in this work, we established that protein coacervates can be used as a platform for the selective enrichment of nanoplastics within droplets. Looking forward, the sequence-level control provided by proteins allows for a large design space to further enhance the coacervate microenvironment, either to further probe the surface features of nanoplastics or to functionalize the droplets for nanoplastic sequestration and degradation.

## Acknowledgements

This work was supported by grants from the National Science Foundation (NSF), National Institutes of Health (NIH), and the Blavatnik Fund for Engineering Innovations in Health. NIH funding was provided under award number R35GM138378, and this material is based upon work supported by the National Science Foundation Graduate Research Fellowship under Grant No. DGE 2437839. Any opinions, findings, and conclusions or recommendations expressed in this material are those of the author and do not necessarily reflect the views of the National Science Foundation. The authors are grateful for the assistance from Dr. Jerry Chang at the Columbia University Precision Biomolecular Core Facility regarding fluorescence spectroscopy techniques and the guidance from Dr. Nico Mendez on nanoplastic formation and characterization from the Kumar Group at Columbia University.

## Supporting information

The supplementary information includes additional data, including amino acid sequences, SDS-PAGE images, and fluorescence excitation and emission spectra.

