## Supplementary Information for "Coacervate droplet sequestration of heterogenous nanoplastics with elastin-like polypeptides"

**Methods**

Unless otherwise noted, all chemicals and supplies were purchased from Fisher Scientific.

**Fluorescent nanoplastic formation and isolation**
Semicrystalline polyethylene terephthalate granules (Goodfellow, ES30-GL-000115), polyamide granules (Nylon 6, Goodfellow, AM30-GL-000122), and polystyrene pellets (Scientific Polymer Products, cat# 844) at 0.4 g plastic/mL ultrapure water (Sigma Aldrich) were stirred at room temperature at 400 rpm for 1 m.[1] For a schematic of the nanoplastic isolation process, reference SI Fig. 1. For non-fluorescent nanoplastic samples of PET, PA, and PS, the supernatant was collected, sonicated in a water bath for 5 min, and centrifuged at 10,000xg for 15 min (Sorvall Legend XTR) in 50 mL polypropylene centrifuge tubes (Corning, 430829) to pellet out macro- and microplastics. The top 90% by volume of the nPET and nPA was carefully transferred to a new tube and centrifuged at 10,000xg for 1 h. The top 80% of the supernatant was then removed and discarded, and the remaining volume was collected and stored in glass vials for characterization. Given the lower density, following the initial centrifugation step, the top 90% by volume of the nPS solution was transferred to a glass centrifuge tube and left to rest undisturbed. After a week, the top 20% was removed and stored in glass vials for characterization.

For fluorescent nanoplastic samples, the supernatant after 1 m of stirring was collected into glass scintillation vials and a 1 mg/mL Nile Red (Sigma Aldrich) stock solution in ethanol was added to achieve a final concentration of 2 µg/mL Nile Red in each vial.[2] The samples were then heated for 3 h in a water bath at 70 °C for PET ($T_g$ = 85–95 °C[3]) and PS ($T_g$ = 85–105 °C[4]), and 40 °C for PA ($T_g$ = 65–80 °C[5]) given its lowered glass transition temperature. After cooling, the solution was sonicated in a water bath for 5 minutes, transferred to 50 mL tubes, and centrifuged at 10,000xg for 15 min to pellet out macro- and microplastics. For PET and PA samples, the top 90% was transferred to a new tube and centrifuged at 10,000xg for 1 h. The top 80% of the supernatant was then removed, and the remaining sample was resuspended to its original volume in ultrapure water. After three subsequent cycles of centrifugation and dilution to remove any remaining free dye, a final spin at 10,000xg for 1 h was used to concentrate the fluorescent nanoplastics and the bottom 20% of the solution was stored in a glass vial. For the dyed PS samples, the same week-long undisturbed settling process was repeated for two cycles. Carboxy-nPS (ThermoFisher, F8801) was diluted 5000-fold into ultrapure water. The particle size distributions of the non-fluorescent and fluorescent nanoplastics was determined by dynamic light scattering (DLS), nanoparticle tracking analysis (NTA), and confirmed via scanning electron microscopy (SEM) as described below.

**Scanning electron microscopy (SEM)**
Silicon wafers were probe sonicated for 10 min in a methanol bath and dried with a stream of nitrogen gas. Nanoplastic solutions were dropcast onto cleaned silicon wafers (1 cm by 1 cm) and left to dry overnight. The samples were attached to stubs with copper tape, sputter coated (2 nm of 5% Pd/Au), and imaged using a Zeiss Sigma VP Scanning Electron Microscope ($6.6 \times 10^{-5}$ mbar, 5.00 kV). Both undyed and fluorescent nPS were dropcast with 50% ethanol to aid in particle dispersion while drying.

**Dynamic light scattering (DLS)**
Dynamic light scattering measurements were performed using a Zetasizer Nano ZS with a 633 nm laser article size distributions and zeta potentials. Nanoplastic stock solutions (70–200 µL) were transferred into ultramicro cuvettes (ZEN0040, BrandTech 759220) for size distribution measurements and 1 mL of each nanoplastic sample was transferred into a dip cell (ZEN1002) for zeta potential measurements. Scattering intensity autocorrelation functions were collected using automatic attenuator selection and a 4.65 mm measurement position with non-invasive backscatter detection at 173°. Three measurements were taken per nanoplastic type, with 10–20 runs each for the size distributions and up to 40 runs for zeta potential. 0.05% v/v TWEEN® 20 (Sigma Life Science) was added to nPS for size measurements to stabilize the smaller particles and prevent aggregation. All other samples were suspensions in the water solutions the nanoplastics were formed in. Zeta potential measurements applied the Smoluchowski approximation, and the stock solutions ranged from pH 6.8–7.2.

**Nanoparticle tracking analysis**

NTA measurements were performed using a Malvern NanoSight NS300 with a red 638 nm laser and a scientific CMOS camera. Samples were sonicated in a water bath for 5 min prior to measurement. Three 100 µL samples from the same batch of nanoplastic were measured in sequence. Two 30 s videos were recorded and analyzed for each sample, and all six runs were averaged for the particle size histogram.

**ELP expression and purification**

The expression and purification protocols of ELP-K30, ELP-DE30 and ELP-V150 were adapted from prior work.[6] All proteins were expressed in NiCo21(DE3) *E. coli* cells. ELP-K30 and ELP-DE30 plasmids were cloned via polymerase cycling assembly, and the ELP-V150 plasmid was obtained from Addgene (pET-24a-ELP[V-150], plasmid # 67015).[7] Large scale cultures, 1 LB with 100 µg ampicillin, were inoculated using 5 mL cultures grown overnight in LB media. Cultures were shaken for 3–4 h at 37 °C until the optical density at 600 nm ($OD_{600}$) reached 0.8–1.0, then induced with 1 mM isopropyl β-D-1-thiogalacto-pyranoside (IPTG) and grown for 16–24 h at 25 °C. Cells were collected via centrifugation at 4,000 rpm for 10 min and resuspended in lysis buffer (20 mM tris, 50 mM–300 mM NaCl, pH 8.0). Cells were lysed in an ice bath with a 1/2" probe sonicator (2 s on, 4 s off, 40% amplitude, elapsed time of 30 min) and centrifuged at 10,000xg for 40 min at 4 °C. The supernatant was decanted and filtered through a 0.22 µm PES membrane, yielding a clarified soluble lysate.

For ELP-V150 and ELP-DE30, the clarified lysate was subjected to inverse transition cycling (ITC) to purify the ELP. Briefly, a 5 M NaCl stock solution in 20 mM tris, pH 8.0 was added to the filtered lysate to achieve a final concentration of 2 M NaCl. For ELP-DE30, the lysate was then adjusted to pH 1.8–2.5 with 1 M HCl. The lysate was heated in a water bath at 40 °C for 30 min, then centrifuged at 10,000xg for 15 min at 40 °C. After the hot spin supernatant containing the soluble impurities was removed, the pellet containing the ELP was resuspended with cold phosphate buffered saline (PBS, 137 mM NaCl, 2.7 mM KCl, 8 mM $Na_2HOP_4$, 2 mM $KH_2PO_4$, pH 8.0) and incubated on ice for 30 min. The solution was centrifuged at 10,000xg for 15 min at 4 °C, and the supernatant containing the solubilized ELP was collected for protein identity confirmation via SDS PAGE. The polypeptide solutions were dialyzed into 10 mM tris, pH 8.0 using a 3.5 kDa regenerated cellulose membrane, quantified using the Pierce™ BCA Protein Assay, and aliquoted for storage at 4 °C or -20 °C.

For ELP-K30, the clarified lysate was purified via FPLC with a 5 mL column (Biorad) hand packed with POROS™ XS Strong Cation Exchange Resin (Thermofisher) using an ÄKTA Start (Cytiva). After sample application, the resin was washed with 100 mL of a 50 mM NaCl wash buffer (in 20 mM tris, pH 7.0) followed by another 50 mL wash of a 100 mM NaCl wash buffer (in 20 mM tris, pH 7.0). The protein was then eluted with increasing concentrations of NaCl using 2 CV fractions of EB (20 mM tris, pH 7.0) with 100 mM NaCl stepwise increases from 200 to 600 mM NaCl, followed by 3 CV of elution buffer with 1 M NaCl. A 100 mM ethylenediaminetetraacetic acid (EDTA) stock solution was added to fractions with the polypeptide of interest to achieve a final concentration of 10 mM EDTA to inhibit protease activity. Fractions were then analyzed by SDS-PAGE and pure fractions were dialyzed into 10 mM Tris, pH 8.0 using a 3.5 kDa regenerated cellulose membrane. Samples were concentrated via centrifugal ultrafiltration using a 10 kDa molecular weight cutoff (Millipore Sigma), quantified with the Pierce™ BCA Protein Assay, filtered with a 0. 22 µm PES membrane, and aliquoted for storage at 4 °C or -20 °C.

**Turbidimetry assays for in vitro phase diagrams**

ELP-V150 and ELP-K30, ELP-DE30 samples (40 µL) for turbidity measurements were made in glass bottom 384 well plates (Cellvis, P96-1.5H-N). Final ELP concentrations were 1.5 µM ELP-V150 for the simple coacervates and a total of 32 µM ELP-K30, ELP-DE30 for the complex coacervates. Samples were prepared by first mixing 10 mM tris buffer and NaCl in the well, then adding in ELP stock solutions and gently pipetting. Absorbance measurements at 350 nm were recorded with an Infinite 200 Pro plate reader (Tecan). For the simple coacervate temperature gradient, samples were allowed to incubate for 15 min at each temperature either within the plate reader or in an external water bath, shaken, and then read. Absorbance values were converted to % transmittance with $\%T = 10^{2-A}$, and turbidity was calculated as 100 - %T.

**Volume fraction image analysis**

Clear round-bottom 96 well plates (Corning 3788) were incubated with 300 μL of 1% w/v Pluronic F127 solution overnight, then emptied and rinsed with $ddH_2O$ six times. Coacervate samples (200 μL) were prepared the same way as the turbidimetry assays. To image the droplets, ELP-V150 and ELP-DE30 aliquots were fluorescently labeled with Alexa Fluor 488 NHS ester (Invitrogen, A20000). Briefly, the concentrated protein stocks (1 mL) were incubated with the dye for 1 h and isolated using NAP-5 columns (Cytiva) as instructed by the manufacturer. For the simple coacervates, droplets were prepared at room temperature in 1 M NaCl, 10 mM tris, pH 8.0 with 20% fluorescent ELP-V150. For the complex coacervates, a positive charge fraction of 0.6 was used in 20 mM NaCl, 10 mM tris, pH 8.0 with 5% ELP-DE30. 30 minutes after mixing, the samples were spun at 1,000xg for 10 min at room temperature. For the simple coacervates, 180 μL of the supernatant was removed for improved droplet contrast in imaging. The well plates were then imaged using the XcitaBlue Conversion Screen with a manual exposure of 3 s (Biorad Gel Doc XR+), thresholded in ImageJ, and analyzed for particle area using circularity requirements of 0.5-1.0 and size maximum of 20 $mm^2$. A 0%, 20%, and 40% v/v glycerol solution with Alexa Fluor 488 free dye was used to generate a standard curve to relate particle area to droplet volume.[8]

**Optical microscopy**

Glass bottom 384 well plates were treated with 1% Pluronic F127 for 1 h and rinsed 6 times with $ddH_2O$ prior to sample preparation. For simple coacervates, nanoplastic stock solutions were first diluted into 1 M NaCl, 10 mM tris, then mixed with 0.5 μM ELP-V150 to form droplets at room temperature. For complex coacervates, nanoplastics were suspended in 20 mM NaCl, 10 mM tris, then mixed with 2.6 μM ELP-K30 and 1.4 μM ELP-DE30, added sequentially. For the samples with preformed, settled droplets, the mixing order was buffer, salt, and ELPs, with the nanoplastic solutions carefully added on top 2 h after droplet formation. Images were acquired over the course of 4 h with a Nikon Eclipse Ti microscope with x100/1.40 NA oil objective using both DIC and a Sedat quad filter cube (mCherry, 555/28 nm excitation, 685/40 nm emission). Exposure time was set to 5 ms exposure for all DIC images and 60, 30, 200, or 10 ms for nPET, nPA, nPS, or nPS-COOH, respectively. Images were processed using Fiji and fluorescence histograms for each nanoplastic type across both droplets were matched to the fluorescence intensity of the medium concentration of nanoplastics in the complex coacervates.

**Fluorescence spectroscopy**

Solutions of Nile red labeled nanoplastics (200 μL) were water bath sonicated for 5 min, then pipetted into a 5x5mm quartz cuvette (FireflySci, Type 4 UV Quartz 5x5x45mm) and diluted with water or ethanol (200 μL) immediately prior to measurement. Fluorescence spectra were recorded using a Horiba QuantaMaster 8075-22 Sepctrofluorometer using a 0.1 s integration time, 5 nm bandwidth and a 1 nm step size. Emission and excitation fluorescence sweeps for each sample were first collected at $\lambda_{ex}$ = 560 nm and $\lambda_{em}$ = 650 nm, and the peak emission and excitation wavelengths were then used for the reported spectra. The excitation wavelengths used were 560 nm for nPET, nPA, and free Nile red dye, 540 nm for nPS, and 580 nm for nPS-COOH.

**Fluorescent nanoplastic quantification**

Nanoplastic stock solutions in ultrapure water were sonicated in a water bath for 5 min and transferred to a glass conical tube (Corning, 9950215) and centrifuged for 2 min at 2,000*xg* to remove any aggregates. The top 90% of the nanoplastic solution was then used to quantify partitioning, and these stocks were prepared daily. Given possible decreases in nPET concentration from this process, 100 ng/mL nPET was defined by the original stock concentration as measured by NTA and fluorescence standard curves were prepared daily. Clear round-bottom 96 well plates (Corning, 3788) were coated with Pluronic F127 as described above. Samples (120 μL) were prepared by first mixing the buffer, salt, and nanoplastics within the well, then adding in the ELPs and pipetting 8–10x to mix. After 30 min–2 h, the well plates were then centrifuged at 1,000*xg* for 10 min to collect the droplets. The top 100 μL was carefully removed to a clear bottom flat black 96 well plate (Corning, 3904) for fluorescence measurements of the supernatant nPET concentration, and the remaining 18-20 μL of the supernatant was removed to isolate the droplet pellet. 10 μL of this was reserved, diluted to 100 μL, and measured. The pellet is then resuspended in 100 μL of chilled 10 mM tris for simple coacervates or 1 M NaCl, 10 mM tris for the complex coacervates, vigorously mixed by pipetting, and transferred to a flat black 96 well plate.

Fluorescence measurements were recorded using a Biotek Cytation 5 plate reader (Agilent). The nPET samples were mixed with 100 µL ethanol prior to fluorescence measurement at $\lambda_{ex/em}$ = 560 / 650 nm with 20 nm bandwidths, and the nPA and nPS-COOH samples were measured without additional solvent at $\lambda_{ex/em}$ = 560 / 630 nm and $\lambda_{ex/em}$ = 580 / 610 nm with 20 nm bandwidths, respectively. For the droplet reuse cycles, the droplet pellet was resuspended in a room temperature solution of concentrated nPET in 10 mM tris and mixed thoroughly. The samples were then supplemented with 1 M NaCl to initiate droplet formation, and the cycle was repeated. The nanoplastic concentration in the dense phase was calculated using the mass captured in the resuspended pellet divided by the estimated droplet volume from prior sections.

**Supplementary table 1 |** Calculations for approximating nanoplastic mass concentrations.

| stock | average measured conc.[1] (particles/mL) | average size[2] (nm) | D10 diameter[3] (nm) | D90 diameter[4] (nm) | reported plastic density[5] (g/mL) | est. mass concentration[6] (ng/mL) |
|---|---|---|---|---|---|---|
| nPET | $4.38 \times 10^8$ | 140 | 86 | 168 | 1.38 | 870 |
| nPA | $4.00 \times 10^8$ | 170 | 75 | 309 | 1.14 | 1200 |
| nPS | $1.60 \times 10^8$ | 110 | 85 | 273 | 1.06 | 120 |
| nPS-COOH | $8.00 \times 10^8$ | 100 | 35 | 85 | 1.06 | 4400 |

[1] Assuming the particles per frame measured via NTA is representative of a 0.05 nL measurement volume
[2] Assuming nanoplastics are monomodal and uniform in diameter
[3] Bottom 10th percentile particle size for the sample population
[4] Top 90th percentile of particle sizes
[5] Assuming the density of nanoplastics is unaltered from bulk plastic density
[6] Assuming all nanoplastics are uniform spheres

**Supplementary table 2 |** Amino acid sequence for elastin-like polypeptides

| Protein | Sequence | MW (kDa) |
|---|---|---|
| ELP-K30 | HHHHHHSSGGCEKT[VPGKG ]$_{30}$VPETEGGGY | 15,637 |
| ELP-DE30 | HHHHHHSSGGCEKTVPGEG[VPGEGVPGDG]$_{14}$VPGEG VPETEGGGY | 15,468 |
| ELP-V150 | HHHHHHSSGGVG[VPGVG]$_{150}$VGVPWP | 62,936 |
| ELP-V30 | HHHHHHSSGGCEKT[VPGVG]$_{30}$VPETEGGGYGWP | 15,106 |

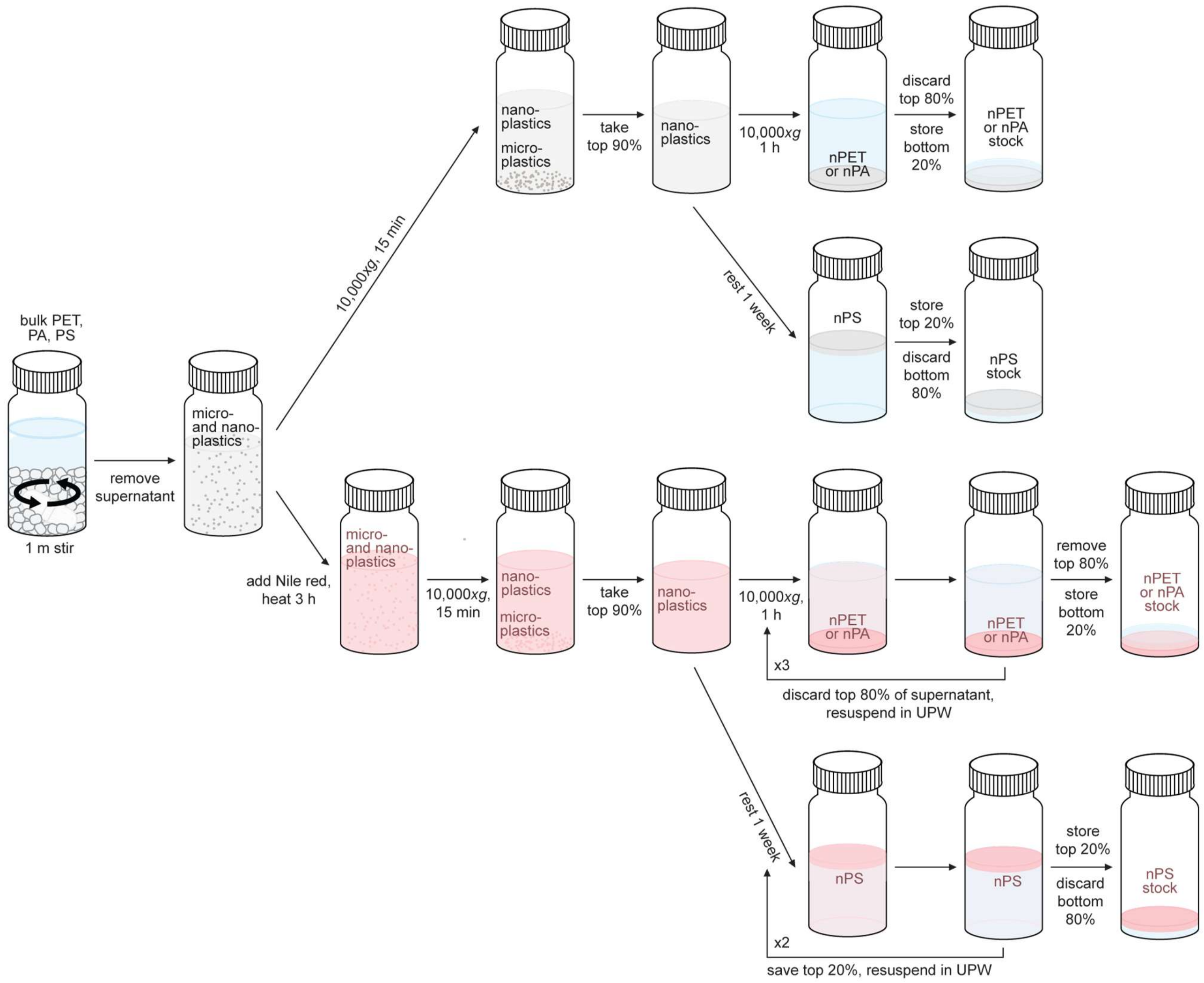

**Supplementary figure 1 | Schematic for fluorescent nanoplastic formation and isolation.** Briefly, bulk plastic pellets are stirred with a PTFE stir bar for 1 m at a concentration of 0.4 g plastic / mL UPW. The solution is then removed and centrifuged to remove the larger microplastics, then centrifuged again to concentrate the nanoplastics. To make fluorescent nanoplastics, Nile red dye (2 μg/mL) is added to the solution and heated for 3 h to facilitate dye diffusion into the micro- and nanoplastics. The dyed solution is then centrifuged to remove the fluorescent microplastics, then concentrated and diluted over several cycles to minimize the concentration of free dye in solution. Centrifugation steps occur in PP conical tubes.

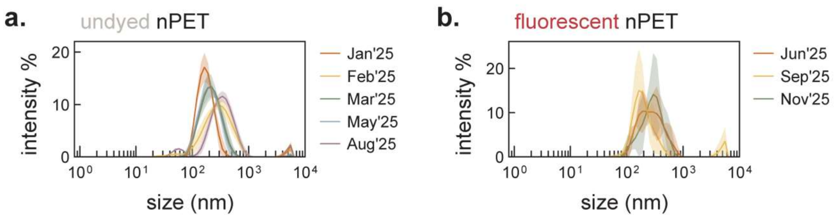

**Supplementary figure 2 | Replicability of nPET production through mechanical abrasion.** Particle size distributions for **a.** independent batches of nPET that was mechanically generated by 1 m of stirring and **b.** independent batches of fluorescently labeled nPET. Data represent the mean value and standard deviation (shaded) of three DLS measurements of each sample.

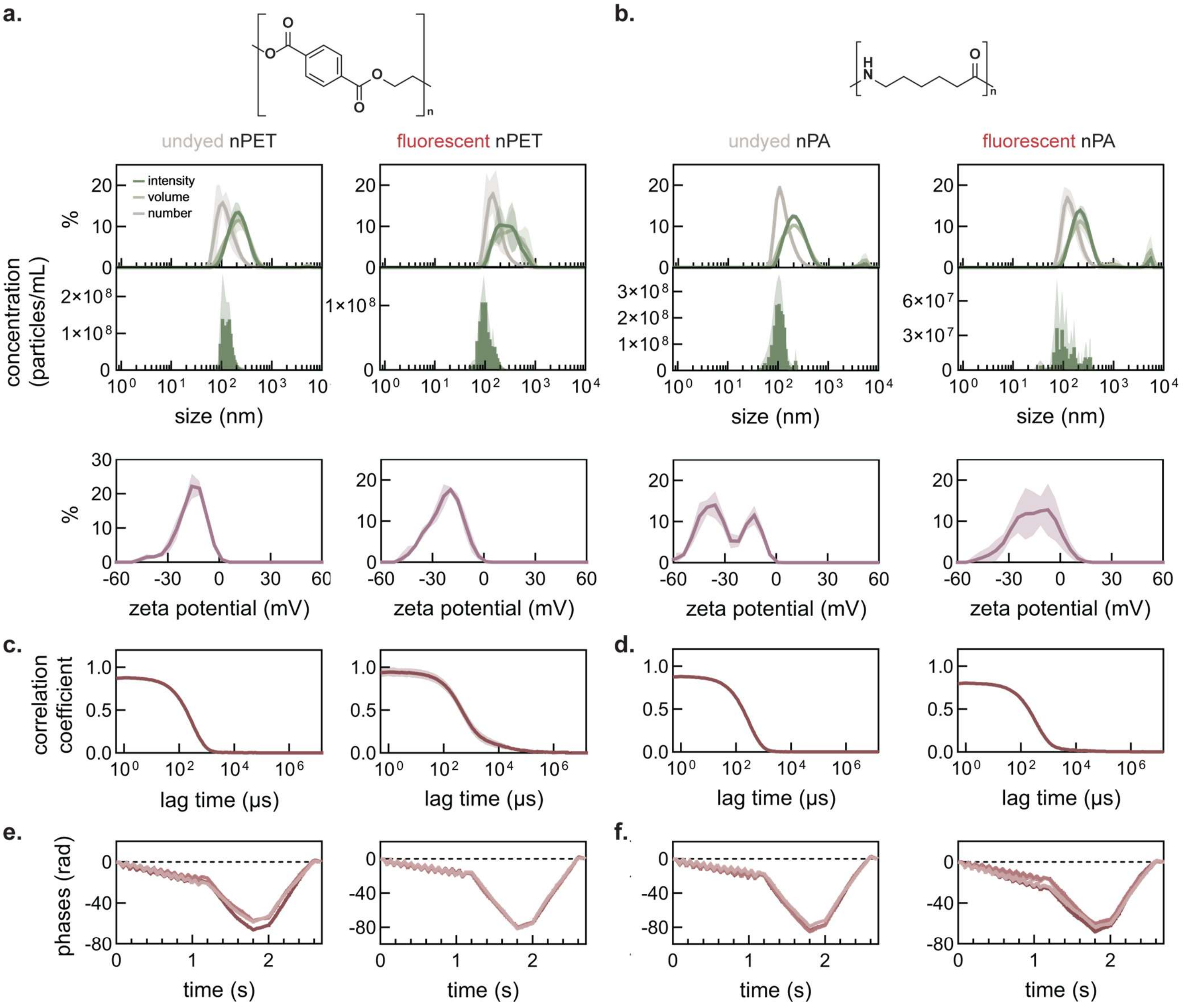

**Supplementary figure 3 | Comparison of PET and PA nanoplastics before and after Nile red labeling.** Particle size distributions from DLS and NTA measurements for **a.** PET and **b.** PA nanoplastics without fluorescent labeling (left) and with Nile red staining (right). Correlograms for DLS size measurements for **c.** nPET and **d.** nPA. Phase plots for zeta potential measurements for **e.** nPET and **f.** nPA. For DLS data, the particle size distribution, zeta potential, and correlogram plots show the mean value and standard deviation (shaded) from three measurements of the same nanoplastic stock solution. Phase plots from each measurement are shown. For NTA data, the histograms represent the mean and standard deviation (shaded) of three technical replicates of the same nanoplastic stock solution.

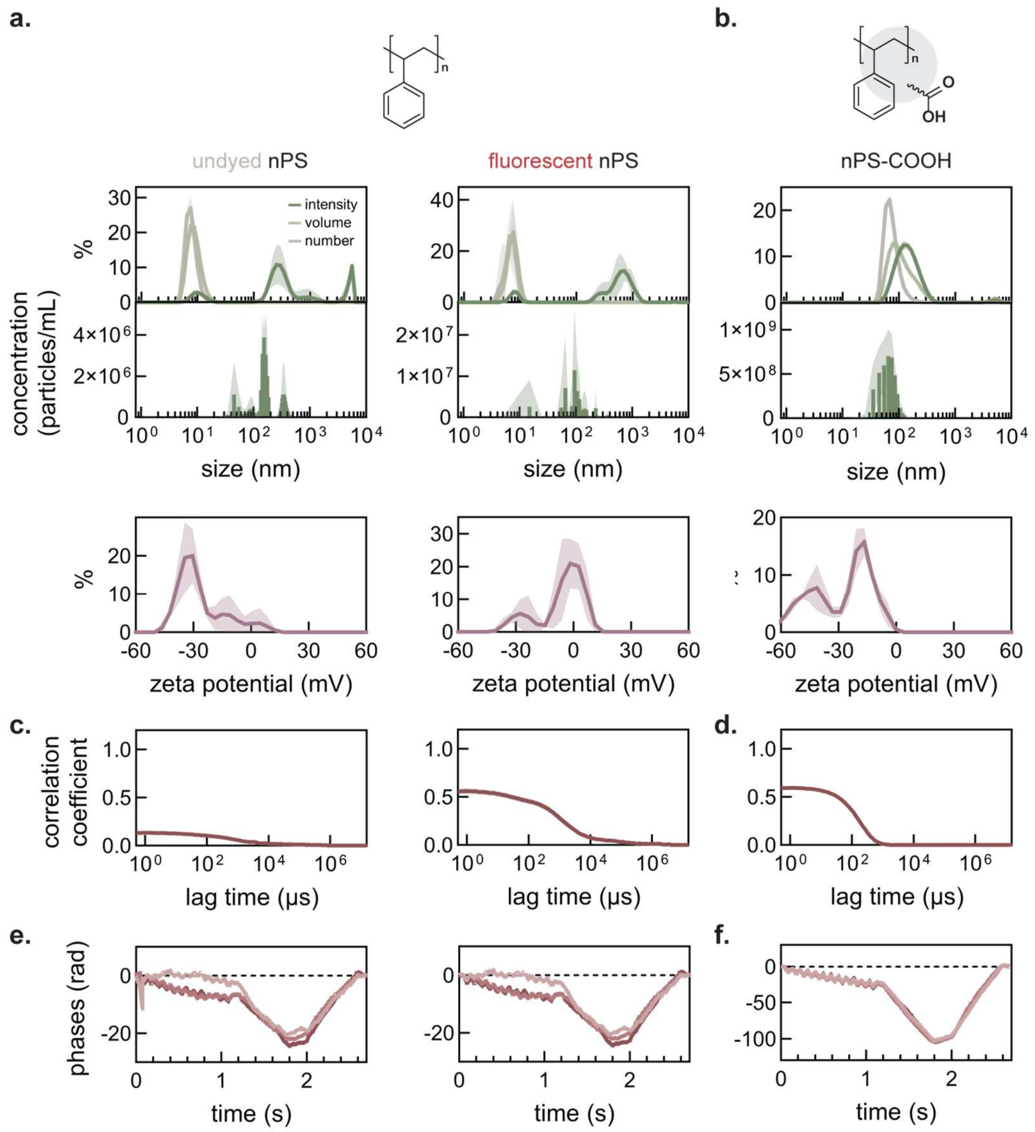

**Supplementary figure 4 | Comparison of PS nanoplastics before and after Nile red labeling.** Particle size distributions from DLS and NTA measurements for **a.** PS without fluorescent labeling (left) and with Nile red staining (right) and and **b.** commercially-made carboxylated latex nanoplastics. Correlograms for DLS size measurements for **c.** nPS and **d.** nPS-COOH. Phase plots for zeta potential measurements for **e.** nPS and **f.** nPS-COOH. Undyed and fluorescent nPS have 0.01% and 0.05% v/v Tween 20, respectively, for DLS particle size distribution measurements to stabilize smaller particles in solution for the duration of measurement. While differences in the zeta potential were observed for undyed and Nile red nPS, we note the poor data quality for undyed nPS can be attributed to an unstable colloidal suspension, as evidenced by the low correlation intercept. For DLS data, the particle size distribution, zeta potential, and correlogram plots show the mean value and standard deviation (shaded) from three measurements of the same nanoplastic stock solution. Phase plots from each measurement are shown. For NTA data, the histograms represent the mean and standard deviation (shaded) of three technical replicates of the same nanoplastic stock solution.

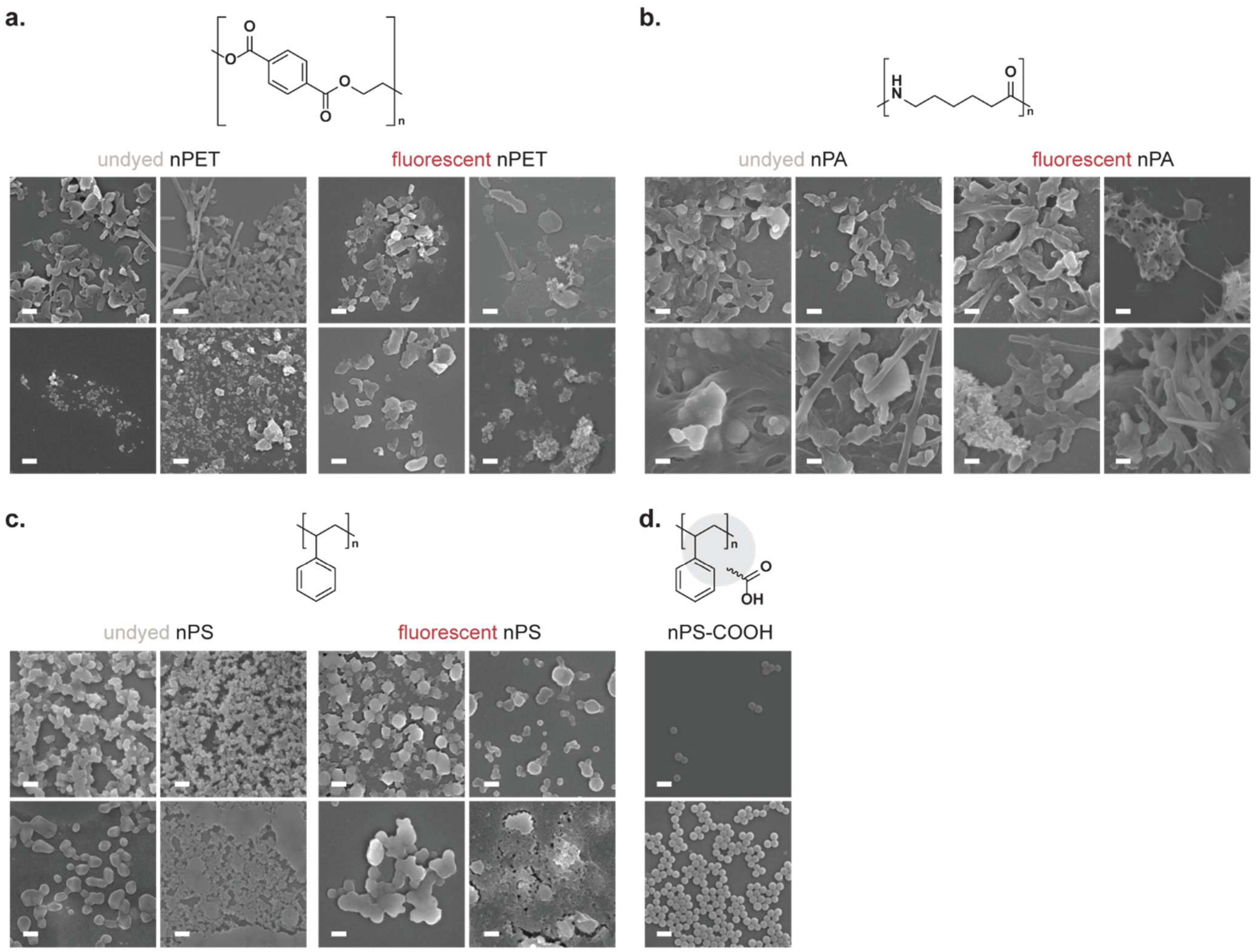

**Supplementary figure 5 | SEM images of unaltered and Nile red labeled nanoplastics.** SEM images of nanoplastics generated through stirring of **a.** PET, **b.** nylon 6 (PA), and **c.** PS in water at ambient conditions for 1 month, and **d.** 0.1 µm carboxylate-modified latex microspheres (nPS-COOH). PS nanoplastics were dropcast with 50% ethanol. The top left image represents the morphology of a majority of the nanoplastics observed across several 10-100 µL dropcast samples, and the additional SEM images highlight other nanoplastics observed in the stock solution. Nanoplastics observed in these images range in size from 20 to 800 nm for the rounder shapes, and 100 nm wide and over several microns long for the fibers. Scale bar, 200 nm.

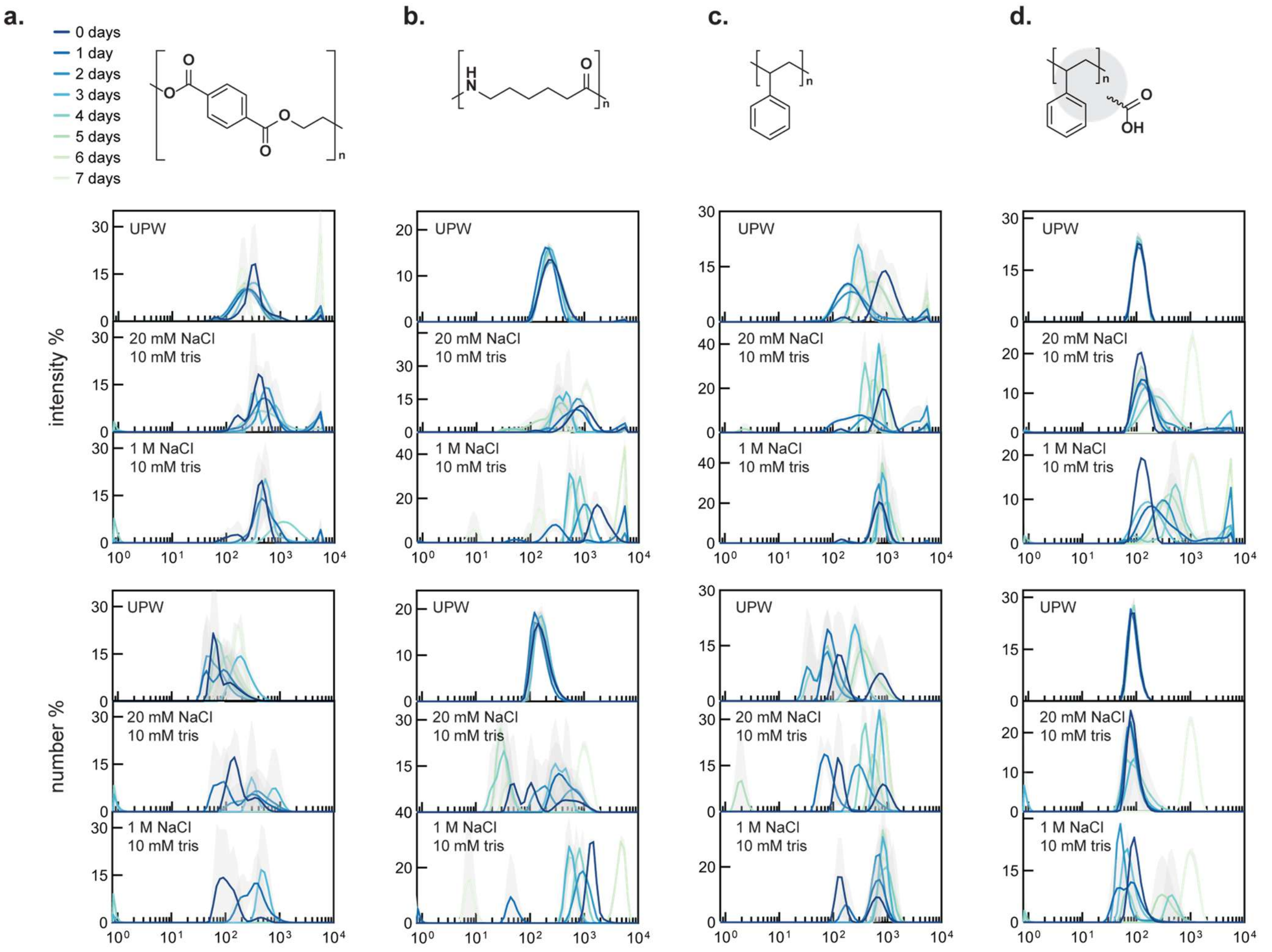

**Supplementary figure 6 | Nanoplastic stability in water, low salt, and high salt solutions.** Intensity (top) and number (bottom) weighted particle size distributions measured by DLS for **a.** nPET, **b.** nPA, **c.** nPS, and **d.** nPS-COOH in varying aqueous solutions (pH ~7, room temperature) over 7 days following an initial sonication of the stock solution and subsequent addition of salts. nPET, nPA, and nPS-COOH in water are relatively stable after a week. No surfactant was added to the nPS samples, so aggregates are observed in initial samples. Increased salt concentrations are shown to destabilize nanoplastic stocks, as we observe larger peaks indicative of particle aggregation for nPET and nPA, while the nPS-COOH remains colloidally suspended for a few days after the addition of salt. Particle size distributions show the mean value and standard deviation (shaded) from three measurements of the same nanoplastic stock solution daily.

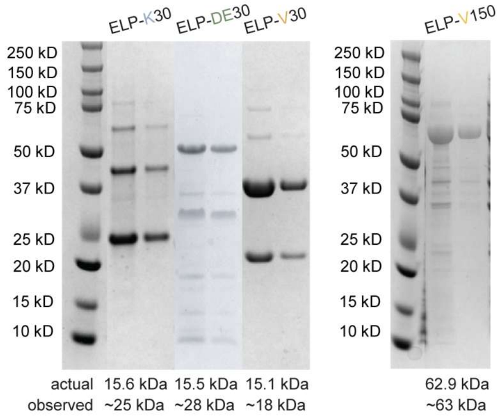

**Supplementary figure 7 | SDS PAGE analysis of purified ELPs.** Protein stock solutions (1–2 mg/mL, approximated by bicinchronic acid assays) of ELP-K30, ELP-DE30, ELP-V30, and ELP-V150 were run on a 4-12% bis-tris gel with NuPAGE LDS loading buffer. Dimers and trimers are present likely due to disulfide bond formation on the N-terminus of the polypeptide, and observed MW of the 30 repeat ELPs are higher than the actual MW of the protein likely due to the disordered and highly charged nature of the protein.

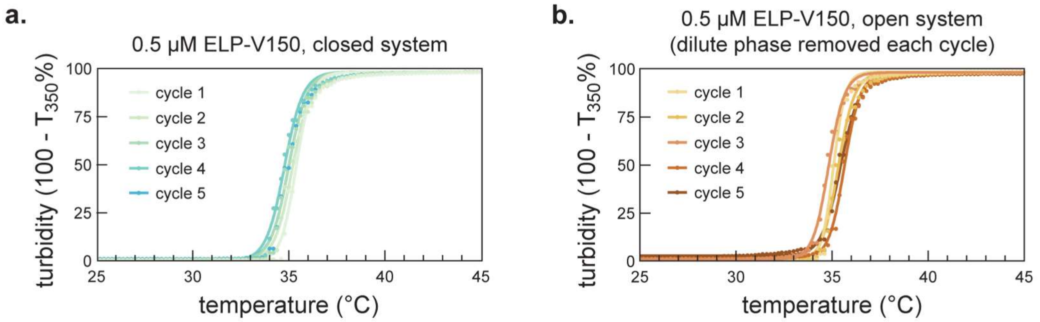

**Supplementary figure 8 | Cloud point measurements of ELP-V150.** Turbidity of a simple coacervate solution of 0.5 μM ELP-V150 in 20 mM NaCl, 10 mM tris, pH 8, measured as a sample is heated at 1 °C/min. **a.** The solution is allowed to reach room temperature and then subjected to the same heating cycle. **b.** The sample is heated to 70 °C and spun down in a heated centrifuge to create a macrophase droplet, and the dilute phase is carefully removed without disturbing the droplet and replaced with chilled, fresh 20 mM NaCl, 10 mM tris solution and subjected to the same heating cycle.

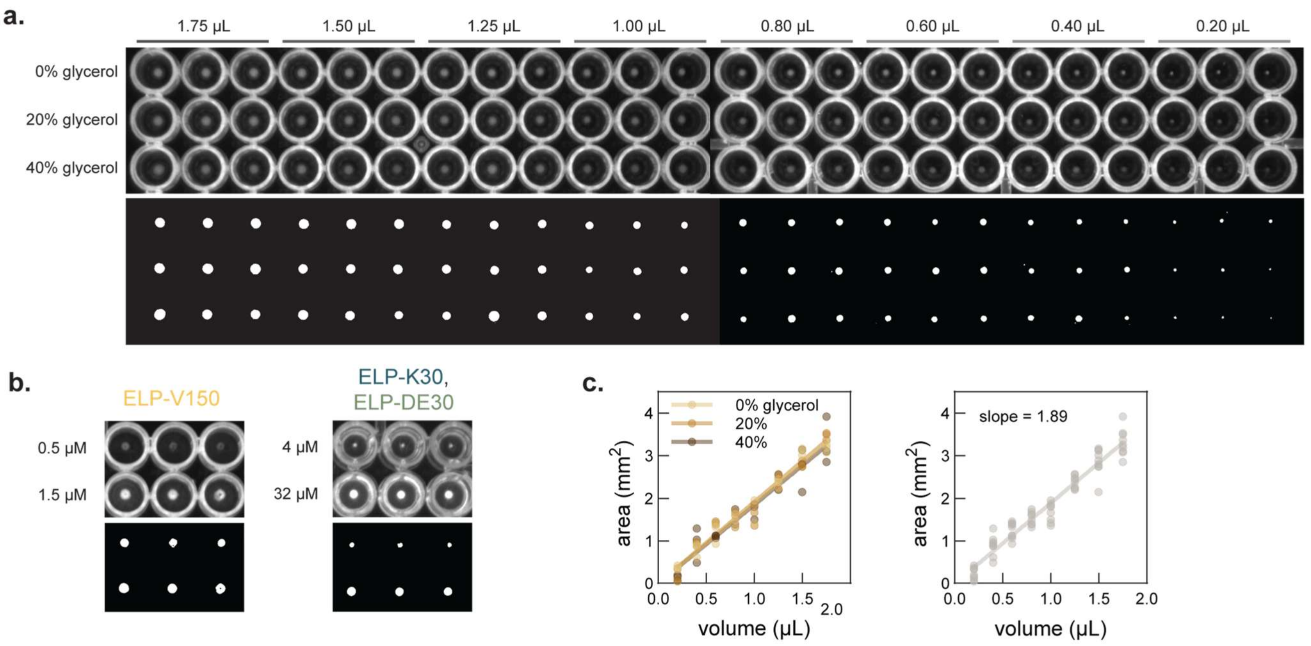

d.

| type | protein | conc. (µM) | volume (µL) | area ($mm^2$) r1 | r2 | r3 |
|---|---|---|---|---|---|---|
| simple | ELP-V150 | 0.5 | 300 | 1.17 | 0.99 | 0.95 |
| simple | ELP-V150 | 1.5 | 200 | 1.60 | 1.44 | 1.49 |
| complex | ELP-K30, ELP-DE30 | 4 | 300 | 0.37 | 0.37 | 0.30 |
| complex | ELP-K30, ELP-DE30 | 32 | 100 | 1.27 | 1.14 | 1.12 |

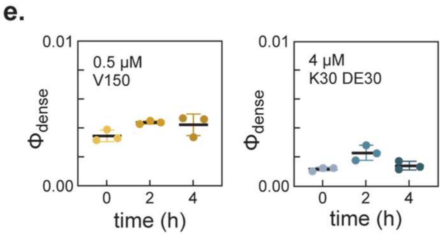

**Supplementary figure 9 | Measurement of simple and complex coacervate volume fractions. a.** Well plate images (top) of glycerol / water stock solutions with approximately 2 µg/mL Alexa Fluor 488 to create standard curves for solutions of varying density. **b.** Well plate images (top) of low and high concentrations of simple and complex coacervates at 1 M NaCl and 20 mM NaCl, respectively. Plates were treated with Pluronic F127 overnight and centrifuged at 1,000xg for 10 min prior to UV trans illumination imaging with an exposure time of 3 s. A mask was applied to the well plate images to remove the well boundaries, then thresholded (bottom) and quantified in ImageJ. **c.** Standard curve for measured area as a function of volume. Solution density does not affect the measured area (left), so the three density solutions were combined for the volume standard curve (right). **d.** Table summarizing experimental setup and measured droplet area for Fig. 2c,f in main text. **e.** Variation in volume fractions at different droplet coalescence times. Volume fraction data points represent technical replicates (n=3), black bar represents the mean, and error bars represent the standard deviation.

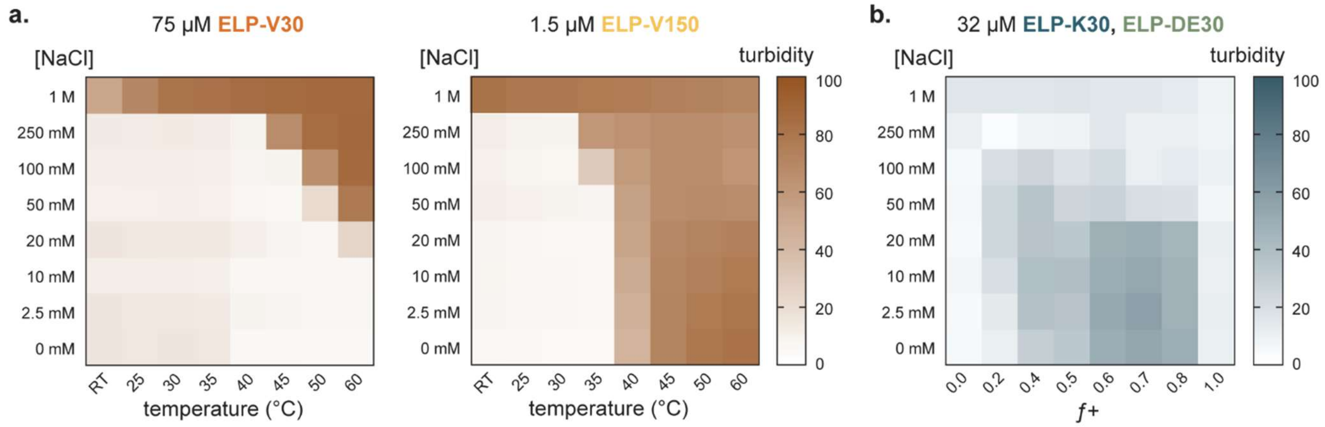

**Supplementary figure 10 | Extended phase diagrams for simple and complex coacervates.** Turbidity (λ = 350 nm) of **a.** 75 μM ELP-V30 and 1.5 μM ELP-V150 samples at varying salt concentrations and temperatures and **b.** total 32 μM ELP-K30, ELP-DE30 at varying charge fractions and salt concentrations. Turbidity heat map values represent the mean of n=3 technical replicates.

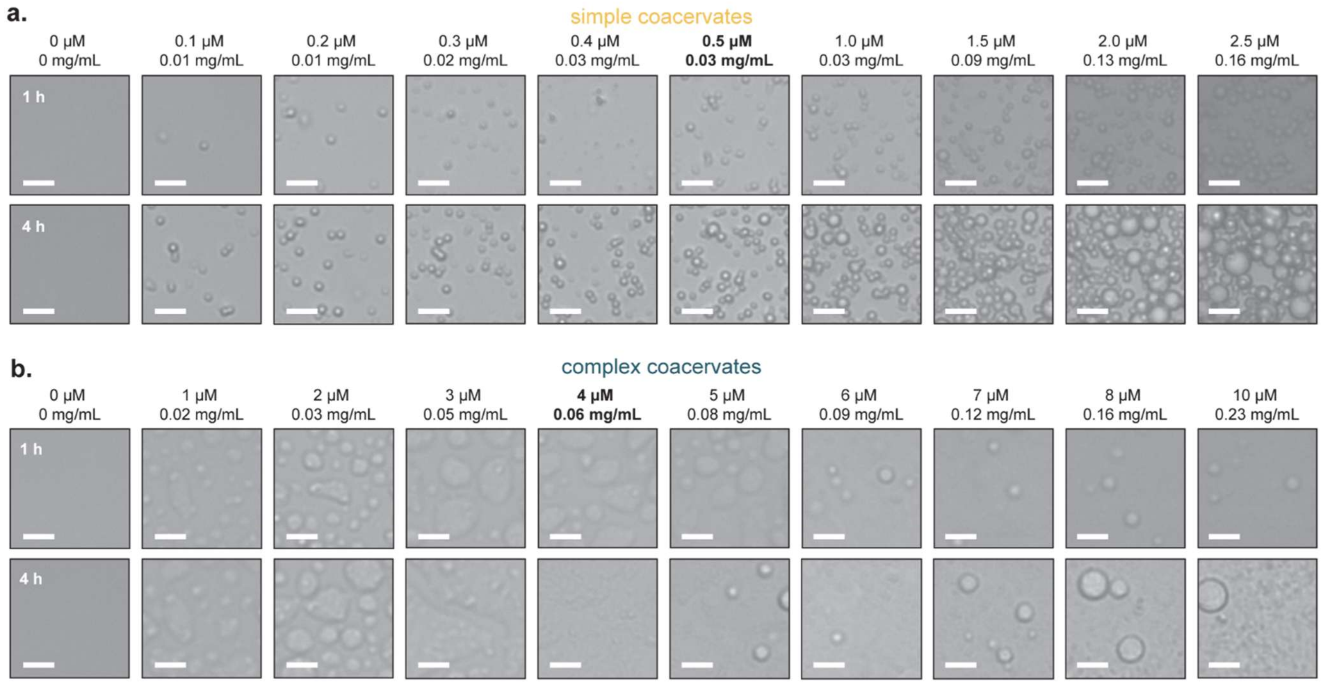

**Supplementary figure 11 | Minimum ELP concentration for droplets observable via microscopy. a.** Brightfield microscopy images of ELP-V150 droplets at 1 M NaCl, 10 mM tris, pH 8.0 at RT over time. **b.** Brightfield microscopy images of ELP-K30, ELP-DE30 droplets at $f+$ 0.6, 20 M NaCl, 10 mM tris, pH 8.0 at RT over time. Scale bar, 10 μm.

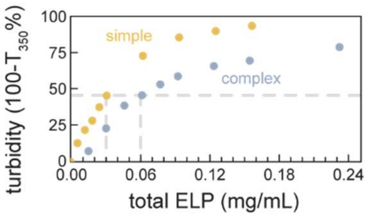

**Supplementary figure 12 | Minimum ELP concentration turbidity measurements.** $A_{350}$ turbidity measured immediately after mixing for samples imaged in SI Figure 10.

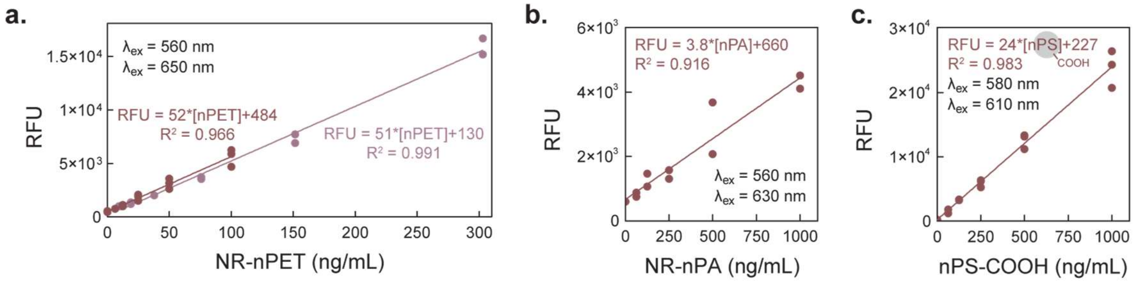

**Supplementary figure 13 | Standard curve for nPET, nPA, and nPS-COOH. a.** Fluorescence measurements of nPET for two sets of serial dilutions (100 ng/mL, n=3, 300 ng/mL, n=2). Samples were diluted in 10 mM tris, and equivolume ethanol was added immediately before measurement. For **b.** nPA (1 µg/mL, n=2) and **c.** nPS-COOH (1 µg/mL, n=3), one set of serial dilutions with 10 mM tris was measured. Individual replicates and a linear regression fit are shown. Excitation and emission wavelenghts selected to maximize the signal to noise ratio. All standard curves were prepared on the day of each sample measurement in a 96 well plate and recorded using a Biotek Cytation 5 plate reader. Data shown represent nPET standard curves recorded for Figure 5 and the nPA and nPS-COOH standard curves recorded for Figure 3g.

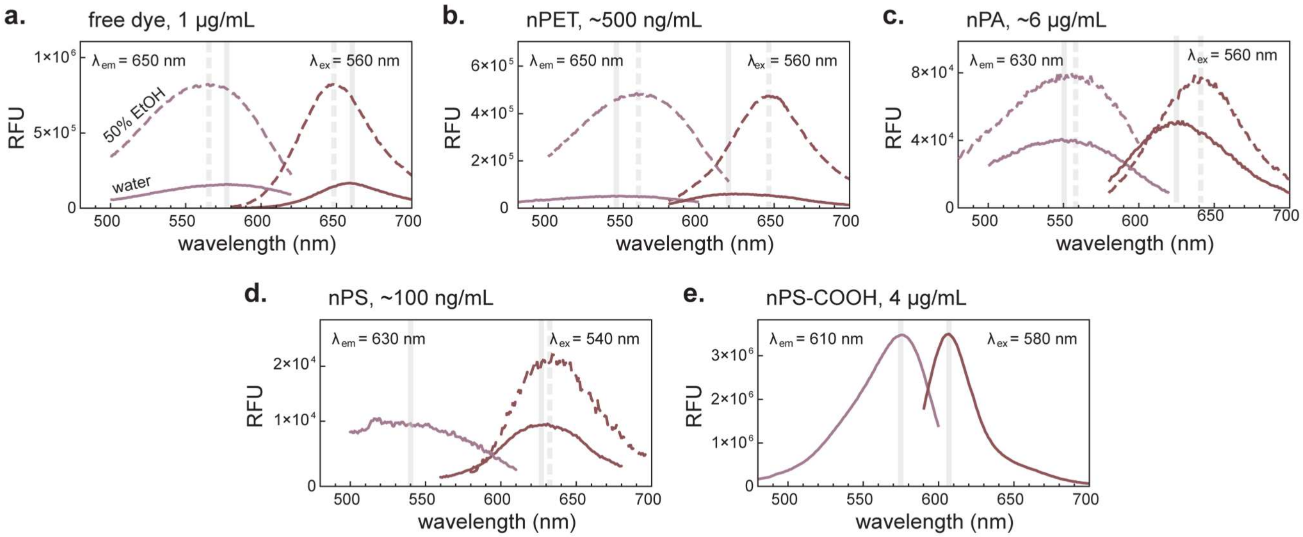

**Supplementary figure 14 | Fluorescence spectra of Nile red dye and labeled nanoplastics.** Peak excitation and emission spectra for **a.** 1 μg/mL Nile red, **b.** ~500 ng/mL nPET, **c.** ~6 μg/mL nPA, **d.** ~100 ng/mL nPS, and **e.** 4 μg/mL nPS-COOH in water (solid line) and 50% ethanol (dashed line) measured with a fluorimeter.

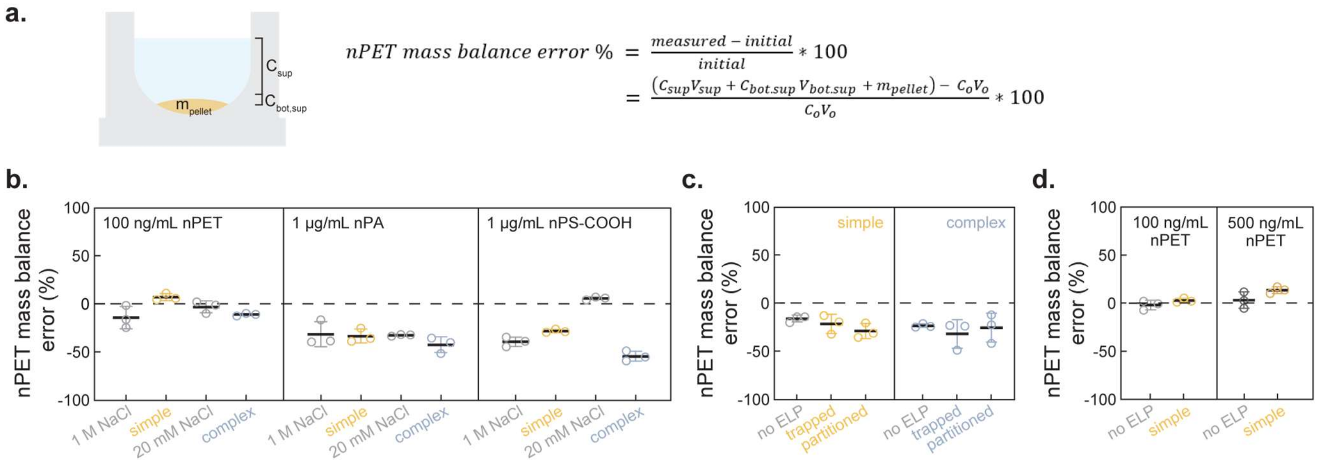

$$nPET\ mass\ balance\ error\ \% = \frac{measured - initial}{initial} * 100$$
$$= \frac{(C_{sup}V_{sup} + C_{bot.sup}\,V_{bot.sup} + m_{pellet}) - C_oV_o}{C_oV_o} * 100$$

**Supplementary figure 15 | Mass balances for quantification of nanoplastic partitioning. a.** Schematic depicting the experimentally measured concentrations and the equation used to calculate the error % to confirm a closed droplet-nanoplastic system and quantify possible nanoplastic loss. Mass balance error for nanoplastic partitioning of **b.** nPET, nPA, and nPS-COOH in Figure 3g, **c.** nPET in Figure 4d, and **d.** nPET in Figure 5b after 3 cycles. Data points represent error % of individual replicates (n=3) and bars represent the mean ± standard deviation of the technical replicates.

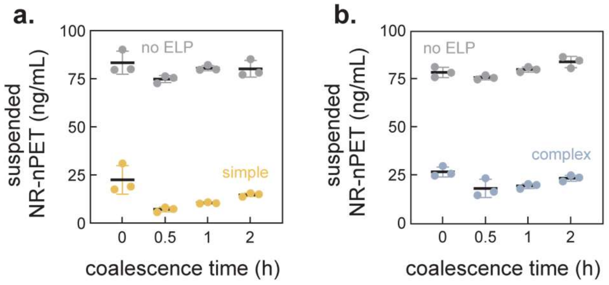

**Supplementary figure 16 | Effect of droplet coalescence time on nanoplastic removal.** nPET solutions with **a.** simple coacervates and **b.** complex coacervates were allowed to rest for varying amounts of time after ELP addition. Droplets were then spun into a macrophase droplet and the nPET remaining in the supernatant was measured. Data represents individual replicates (n=3), bars represent the mean and standard deviation.

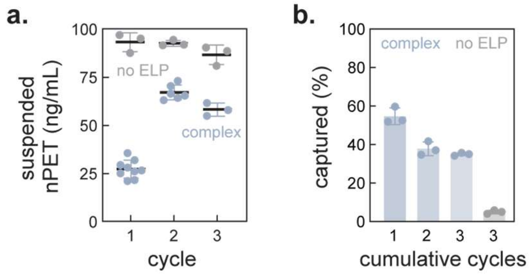

**Supplementary figure 17 | Reusability of complex coacervates for nanoplastic removal.** Using an initial input of 4 µM ELP-K30, ELP-DE30 in 20 mM NaCl at room temperature, the droplets were recovered, dissolved, and exposed to 3 cycles of fresh 100 ng/mL nPET feed solutions. **a.** nPET remaining in the supernatant was measured. Bars represent mean and standard deviation of technical replicates every cycle (n=9 for cycle 1, n=6 for cycle 2, n=3 for cycle 3 of ELP reuse, and n=3 for no ELP control). **b.** Macrophase droplets after 1, 2, and 3 cycles of complex coacervate reuse were dissolved and measured to quantify cumulative nanoplastic capture. Bars represent mean and standard deviation of technical replicates of the reused droplets (n=3 for all cycle counts and no ELP control).

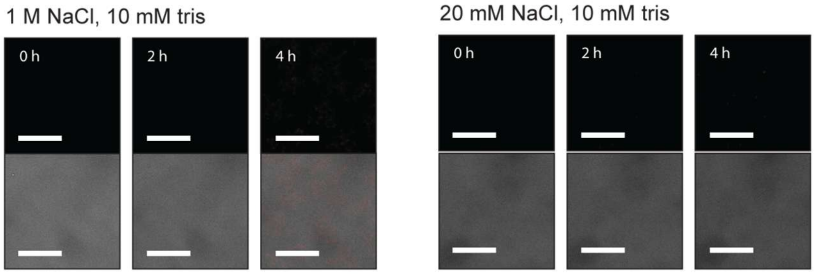

**Supplementary figure 18 | Microscopy of nPET in low and high salt solutions.** 100 ng/mL nPET samples in 1 M and 20 mM NaCl were imaged over 4 h as controls to the partitioning mechanism experiment in Figure 4b-g. No nPET particles were observed to have settled over this time scale. Scale bar, 10 µm.

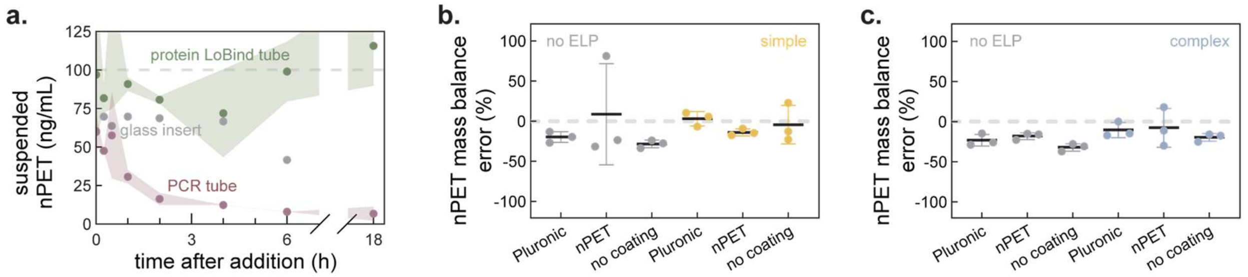

**Supplementary figure 19 | Adsorption of nPET to experiment vessels.** 100 ng/mL nPET solutions were added to **a.** varying tubes with different surface properties to measure surface adsorption. Data points represent the mean and shaded areas represent the standard deviation of the technical replicates (n=3 for protein LoBind tubes and PCR tubes, n=1 for glass vial inserts). For quantification experiments in clear roundbottom well plates, a polymer coating (Pluronic F127) and undyed nPET were used to explore methods to prevent nPET adsorption to the well plates. 100 ng/mL nPET samples in **b.** high and **c.** low salt conditions with and without droplets, were tested to measure nPET loss to adsorption. Data points represent the mass balance error for individual replicates (n=3) and bars represent the mean and standard deviation.
